# Impaired ZNF560 repression during induced pluripotent stem cell reprogramming indicates early epigenetic alterations in Schizophrenia

**DOI:** 10.64898/2026.08.14.744920

**Authors:** Bárbara S. Casas, Elvis Acevedo, Michel Maluenda, Rodrigo Celis, Claudio Letelier-Naritelli, Víctor Pola-Véliz, Stevens K. Rehen, Verónica Palma, Martin Montecino

## Abstract

Despite extensive epigenetic reprogramming, induced pluripotent stem cells (iPSC) from schizophrenia patients (SZ) retain several molecular and functional features of this disease. Transcriptomic and epigenomic analyses were performed in iPSC of SZ and healthy control subjects (HC). Transcriptional profiles were largely similar between SZ and HC iPSC, whereas pronounced differences emerged following neural differentiation, with SZ NSC exhibiting dysregulation of genes involved in neurodevelopment and synaptic function. *ZNF560* was identified as a uniquely and consistently upregulated gene in SZ iPSC, robustly discriminating SZ from HC iPSC. Epigenomic profiling revealed increased chromatin accessibility and reduced DNA methylation at the *ZNF560* promoter in SZ iPSC. ChIP-seq data suggested that ZNF560 can bind to promoters of genes implicated in synaptic signaling and neuronal development. Moreover, a subset of these genes was found to be differentially expressed in SZ neural stem cells. Together, our results identify *ZNF560* as a reprogramming-resistant epigenetic marker of schizophrenia and suggest an altered KRAB-ZNF–mediated regulation in early neurodevelopmental pathways underlying this disorder.

## Introduction

Schizophrenia is a polygenic complex psychiatric disorder with up to 80% heritability. Disease risk has been associated with numerous genetic variants, that can only partially explain the onset and/or progression of this disorder. The latest GWAS for schizophrenia, indicated that the polygenic risk score explained a median of 0.073 of the variance in liability to schizophrenia [1–3]. Hence, although the genetic component can play a significant role, environmental factors may also contribute to the risk by interacting with the different genetic predispositions [4, 5]. Given the significant influence of environmental exposures, the epigenetic regulation of gene expression has a central focus of schizophrenia research. Accordingly, abnormal DNA methylation and altered expression of long non-coding RNAs have been associated with schizophrenia symptomatology [6, 7].

Physio-pathologically, schizophrenia has been associated with synaptic alterations, including reduced synaptic density, down-regulated expression of synapse-associated proteins, and decreased glutamatergic/GABAergic function [3, 8, 9]. Due to the inability of accessing live human brain tissue, we and others have relied on the use of induced pluripotent stem cells (iPSC) to model early stages of this condition [10]. iPSC-derived neural lineage cells obtained from schizophrenia patients (SZ) exhibit phenotypic and molecular features that are characteristic of the disease [11, 12]. Schizophrenia-derived neural progenitors and neurons display synaptic abnormalities, reduced neuronal connectivity, impaired migration, increased oxidative stress, and mitochondrial dysfunction, which were reported in the brains of patients. Additionally, brain organoids generated from schizophrenia iPSC revealed defects in proper neuronal stratification, migration, and connectivity [13, 14]. At the cellular and molecular level, alterations have also been associated with changes in neuronal patterning, proliferation and migration capabilities, synapse formation, oxidative stress, and immune responsiveness [15–20]. Importantly, many of these altered processes overlap with differentially methylated patterns at gene regulatory sequences observed both in *postmortem* brain samples and in peripheral cells from SZ [21, 22]. This finding is particularly relevant, as epigenetic marking is thought to be largely reset during reprogramming and differentiation, hence providing evidence to propose phenotypic convergence phenomena that reflect the emergence of disease-specific features during differentiation. However, whether these epigenetic features are already detectable at the earliest stages of neural commitment, or even prior to differentiation, remains unknown.

Here, we performed comprehensive transcriptomic and epigenomic analyses on iPSC derived from healthy control subjects (HC) and SZ, alongside their neural stem cell (NSC) derivatives to determine whether disease associated molecular signatures emerge at this early developmental stage. Our results indicate that early transcriptomic and epigenetic differences are found in schizophrenia-derived iPSC respect to HC samples. Importantly, we propose that these early alterations here identified may serve as a basis for developing novel disease biomarkers.

## Materials and methods

### Cell lines

Detailed information on patients and the source of cells is provided in Supplementary Table 1. We selected SZ iPSC that came from patients with known familiar risk of SZ, assuring a high genetic component. Culture conditions for iPSC and isogenic fibroblasts are described in Supplemental Methods.

Four iPSC lines (two from HC subjects and two from SZ) were differentiated into NSC following previously described protocols [18].

### RNA-seq

1 µg of total RNA was sent to Quick Biology (https://www.quickbiology.com/; Monrovia, CA, USA) for a standard RNA sequencing service; PolyA ARN, 30M reads per sample, paired-end.

Publicly available datasets used in this article are listed in Supplementary Table 3.

### ATAC-seq

6*10^5^ cells were cryopreserved in mTeSR1 with 5% DMSO and sent to Quick Biology for a standard ATAC-seq service, 50M reads, paired-end.

### Bioinformatic analysis

RNA-seq and ATAC-seq processing, integration of GWAS and ATAC-seq data, Motif enrichment analysis and FIMO analysis are described in Supplemental Methods.

### Methylation sequencing

To identify 5mC methylation in genomic DNA samples, 500 ng of DNA was bisulfite converted using EZ DNA Methylation-Gold Kit (Zymo Research Cat. No D5005, Irvine, CA). Regions of interest were amplified with converted primers and analyzed using BiQ Analyzer [27]. Detailed methods are described in Supplemental Methods.

### Chromatin immunoprecipitation (ChIP)-PCR

ChIP was performed as described previously [28, 29], with minor modifications. Detailed protocol is described in Supplemental Methods.

### Data availability

RNA-seq datasets have been deposited in Gene Expression Omnibus GSE325013 and ATAC-seq data in GSE325014.

## Results

### Differentially expressed genes in iPSC from SZ

To determine transcriptional differences in iPSC derived from schizophrenia patients (SZ iPSC) that reflect early changes in gene expression following cell reprograming, RNA-seq analysis was performed using iPSC samples from three control subjects (HC iPSC) and four SZ. Additionally, these transcriptomic analyses were performed in four iPSC lines (two HC and two SZ) that had been induced to differentiate to NSC. Principal component analysis (PCA) of these transcriptomes revealed distinctively separated populations of transcripts between iPSC and NSC (Supplementary Figure 1A). As expected, iPSC highly expressed canonical pluripotency markers and barely exhibited expression of neural lineage genes, whereas NSC expressed well-characterized early neural lineage commitment genes and showed reduced pluripotency marker expression (Supplementary Figure 1B).

The overall transcriptome of both HC and SZ iPSC showed very similar transcript levels (Figure 1A). Significant (2fold or more, p < 0.05) differences in transcript abundance mostly arose upon neural differentiation engagement (Figure 1B), including 1068 differentially expressed genes (DEGs) in schizophrenia NSC. Gene ontology (GO) analysis of the upregulated DEGs in SZ NSC revealed enrichment of biological processes that included neural system development, neurogenesis and synaptic function (Figure 1C). Interestingly, among the enriched terms we identified glutamatergic and dopaminergic synapse and nicotine addiction (Figure 1D), which have been reported as significantly associated with the schizophrenia symptomatology [9]. On the other hand, GO analysis of the downregulated DEGs in SZ NSC showed terms related to developmental processes, including cell differentiation (Figure 1C). Moreover, among the enriched altered pathways identified were PI3K-AKT and ECM signaling, which also have been recently linked to schizophrenia [30–33].

**Figure 1.**
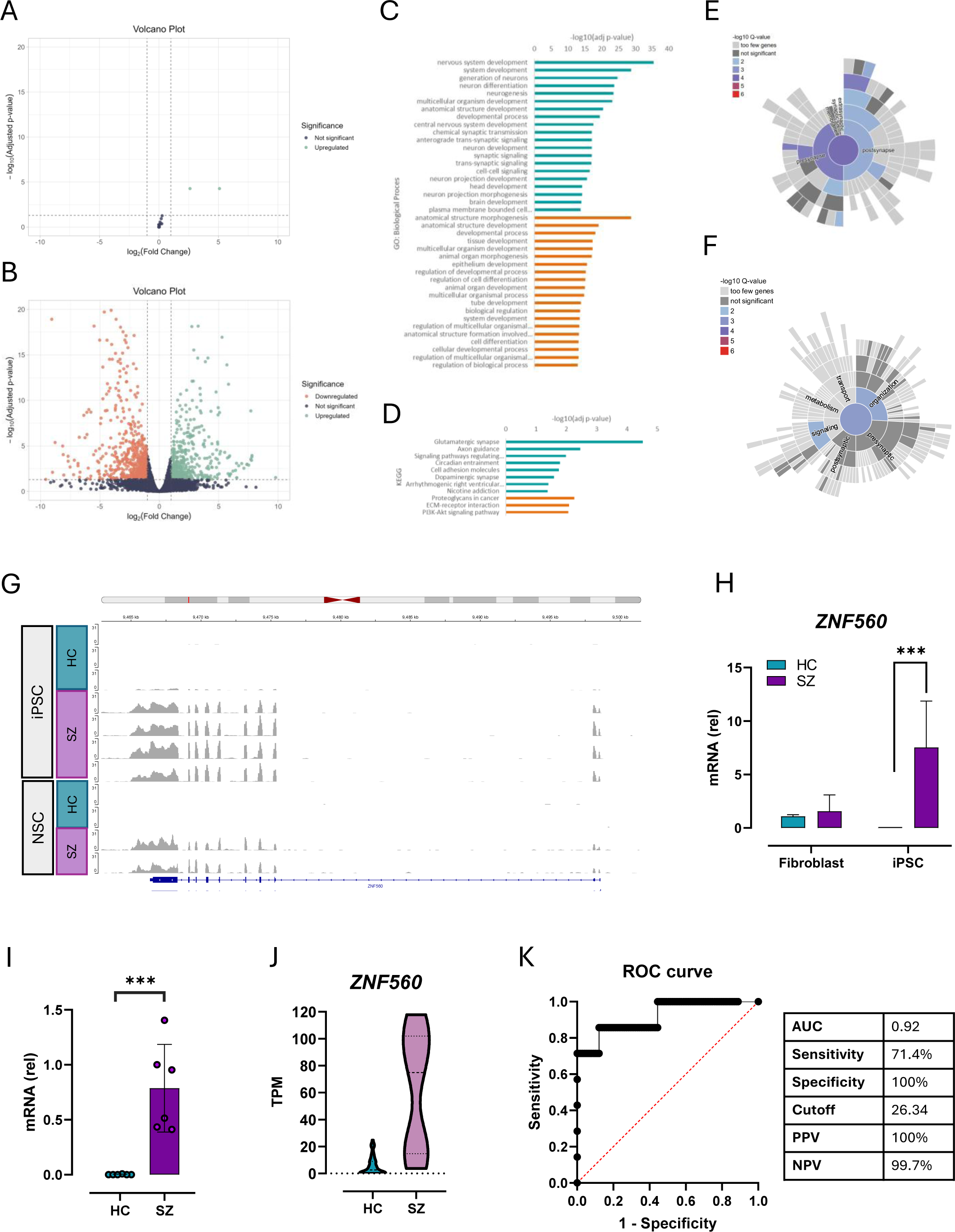
Transcriptional analysis of HC and SZ iPSC. **A-B.** Volcano plot showing a differential expression analysis performed on RNA-seq data from iPSC cultures (3 HC and 4 SZ) (A) and NSC cultures (2 HC and 4 SZ) (B). Genes with an adjusted p-value < 0.05 and |log2 fold change| > 1 were considered differentially expressed. **C.** GO analysis of biological processes of NSC RNA-seq data. The top 20 terms are shown in green (for upregulated genes in SZ NSC) or orange (for downregulated genes in SZ NSC). **D.** GO analysis of biological pathways. Terms are shown in green (for upregulated genes in SZ NSC) or orange (for downregulated genes in SZ NSC). **E-F.** Sunburst diagram of significantly enriched ontology terms for location (E) or function (F) obtained by SynGO analysis of NSC DEGs. **G.** Visualization of tracks with RNA-seq reads at ZNF560 locus on chromosome 19. Data for all iPSC and NSC samples is group-autoscaled using Integrative Genome Viewer (IGV). **H.** mRNA levels of ZNF560 assessed by RT-qPCR in iPSC and the isogenic fibroblasts from which they were originally reprogrammed; *GAPDH* was used as a housekeeping gene. Data are shown as mean ± SD, relative to Fibroblast HC1, with *** p ≤ 0.001 by 2-way ANOVA. **I.** mRNA levels of ZNF560 assessed by RT-qPCR in 6 HC and 6 SZ iPSC; *GAPDH* mRNA was used as a normalizing expression control. Data are shown as mean ± SD, relative to iPSC SZ1, with *** p ≤ 0.001 by t-test. **J.** TPM values of ZNF560 for HC-hiPSC and SZ-hiPSC (n=107). The graph includes all datasets from Supplementary Table 3. **K.** ROC analysis of data from J.

To further characterize synaptic-associated transcriptional alterations in SZ NSC, we performed a SynGO analysis [34]. Among the 1068 DEGs, it was found that 135 mapped to SynGO annotated genes. This analysis identified 12 cellular components and 3 biological processes as significantly enriched (FDR <0.01), corresponding to proteins localized mainly at the pre- and post-synapse, including presynaptic membrane and postsynaptic structural components, with significant enrichments for specialized postsynaptic density (Figure 1E, Supplementary Table 4) and trans-synaptic signaling terms (Figure 1F, Supplementary Table 4).

We next searched for overlaps between DEGs identified in SZ NSC and DEGs reported in *postmortem* dorsolateral prefrontal cortex (DLPC) of schizophrenia patients found at publicly available datasets of the CommonMind Consortium initiative. This dataset includes samples from 258 schizophrenia patients and 279 control subjects (SZDB 2.0 database) [35]. We found that 33 DEGs in SZ NSC were also similarly affected in the DLPC of schizophrenia patients, including genes involved in synapse function, such as *CALB1*, *CPLX1,* and *GPRIN3* (Supplementary Table 5). Together, these results indicated that following induction of neural commitment, SZ NSC iPSC exhibited an expression profile distinct from that of control NSC, including a subset of DEGs comparable to those observed in the cortex of SZ.

On the other hand, despite this overall similarity in transcript expression between HC and SZ iPSC (Figure 1A), two DEGs were identified: *ZNF560* and *HERC2P3* (Figure 1A). *ZNF560* was found to be significantly expressed in SZ iPSC samples and, in contrast, was below the level of significant detection in HC iPSC (Figure 1G). This pattern of differential *ZNF560* mRNA expression between SZ and HC samples was maintained following the commitment of both iPSC types to NSC. By contrast, *HERC2P3* mRNA expression is further enhanced as SZ iPSC engage neural differentiation, whereas HC iPSC exhibited comparably lower *HERC2P3* expression in both iPSC and NSC (Supplementary Figure 1C).

We next determined whether this differential *ZNF560* transcript expression was already established in fibroblast samples from which the iPSC were generated. RT-qPCR analysis indicated that *ZNF560* transcript was significantly expressed in the fibroblasts of the control subjects and that this *ZNF560* expression was silenced upon their reprogramming to iPSC. Interestingly, fibroblasts from SZ patients also expressed the *ZNF560* transcript at levels that were comparable to control subjects, and as these SZ fibroblasts were reprogrammed to iPSC, this *ZNF560* mRNA expression remained significantly upregulated (Fig. 1H).

To rule out a reprogramming-method -dependent effect, we assessed *ZNF560* expression in six HC-iPSC and six SZ-iPSC lines; half of them (3 HC and 3 SZ) were reprogrammed using integrating vectors, and the other half through episomal vector delivery. The results confirmed that, regardless of the gene delivery strategy utilized for reprogramming (Figure 1I), *ZNF560* silencing occurred exclusively in HC-derived iPSC, remaining highly expressed in cells derived from SZ patients. To rule out a passage-dependent effect, we confirmed that early passages (p14) of HC-iPSC already lacked *ZNF560* gene expression, and that SZ-iPSC maintained significant *ZNF560* transcript levels up to passage 45 (Supplementary Figure 1C). Together, these results indicated that *ZNF560* expression in SZ iPSC is not a transient “epigenetic memory” of the SZ fibroblast state that should fade out with passaging but rather suggests an intrinsic mechanism that prevents *ZNF560* repression during reprogramming of SZ-derived cells.

We further validated these findings by analyzing publicly available RNA-seq datasets from HC and SZ-iPSC. The true pluripotent condition of each of the iPSC included in the analysis was assessed by confirming the elevated expression of two well-established pluripotency gene markers, *NANOG* and *OCT4* (we established a minimal expression threshold for *NANOG* and *OCT4* >1 TPM for each). 100 HC and 7 SZ iPSC datasets met this pluripotency criterion and were finally considered in the analysis (Supplementary Table 3). *ZNF5c0* expression differed significantly between HC and SZ-iPSC (Figure 1J). ROC analysis of *ZNF560* expression data was then used to assess whether this parameter could be used to efficiently distinguish between the HC and SZ-iPSC groups. ROC curve showed an AUC of 0.92, which classifies *ZNF5c0* expression level in iPSC as a strong discriminator between healthy vs schizophrenia cell samples (Figure 1K). Given the imbalance between groups (100 vs 7), we additionally generated a precision-recall (PR) curve, which has been shown to perform better when evaluating unbalanced data. The PR-AUC was 0.78, more than tenfold above the expected baseline value (0.071), hence confirming that *ZNF560* transcript expression allows good discrimination between HC and SZ-iPSC (Supplementary Figure 1D).

### Differentially accessible chromatin regions in iPSC from SZ

*ZNF5c0* gene is expressed in fibroblasts and subsequently silenced during reprogramming to HC iPSC, an event that fails to occur in SZ iPSC, resulting in persistent *ZNF5c0* expression. Chromatin remodeling and epigenetic regulation are well-established mechanisms controlling gene transcription [36, 37]. Therefore, we hypothesized that an altered epigenetic process at the *ZNF5c0* locus prevents its transcriptional downregulation during the reprogramming of fibroblasts from SZ. To test this hypothesis, we performed chromatin accessibility analysis by ATAC-seq, seeking to determine whether changes in chromatin compaction at the *ZNF5c0* gene were impaired in iPSC from SZ patients with respect to HC.

Comparison of the global accessibility maps between iPSC samples, did not indicate extensive changes in chromatin organization at gene transcriptional start sites (TSS) (Figure 2A). However, as we screened the chromatin accessibility data to identify differentially accessible regions (DARs) between HC and SZ iPSC, we determined that 89 DARs (FDR<0.05) were enriched in SZ iPSC versus HC iPSC, of which 58 DARs showed increased chromatin accessibility in SZ compared to HC iPSC (Figure 2B). Also, several DARs were mapped at distal intergenic genomic regions, as well as DARs reflecting increased accessibility at gene introns in SZ iPSC (about 72%) (Figure 2C-D). To explore the potential functional role of the sequences associated with these DARs, we intersected these genomic loci with data in recent GWAS reporting genomic linkage to schizophrenia [2]. Of 287 loci included in these GWAS, four overlapped with DARs identified in SZ iPSC: one corresponding to a loss in accessibility in SZ iPSC (exon 12 of *USP6* gene) and three corresponding to increased accessibility in SZ-iPSC; two at the *CYTH1* and *CASP12* genes, and one at the promoter of the *H4C6* gene (Supplementary Table 6).

**Figure 2.**
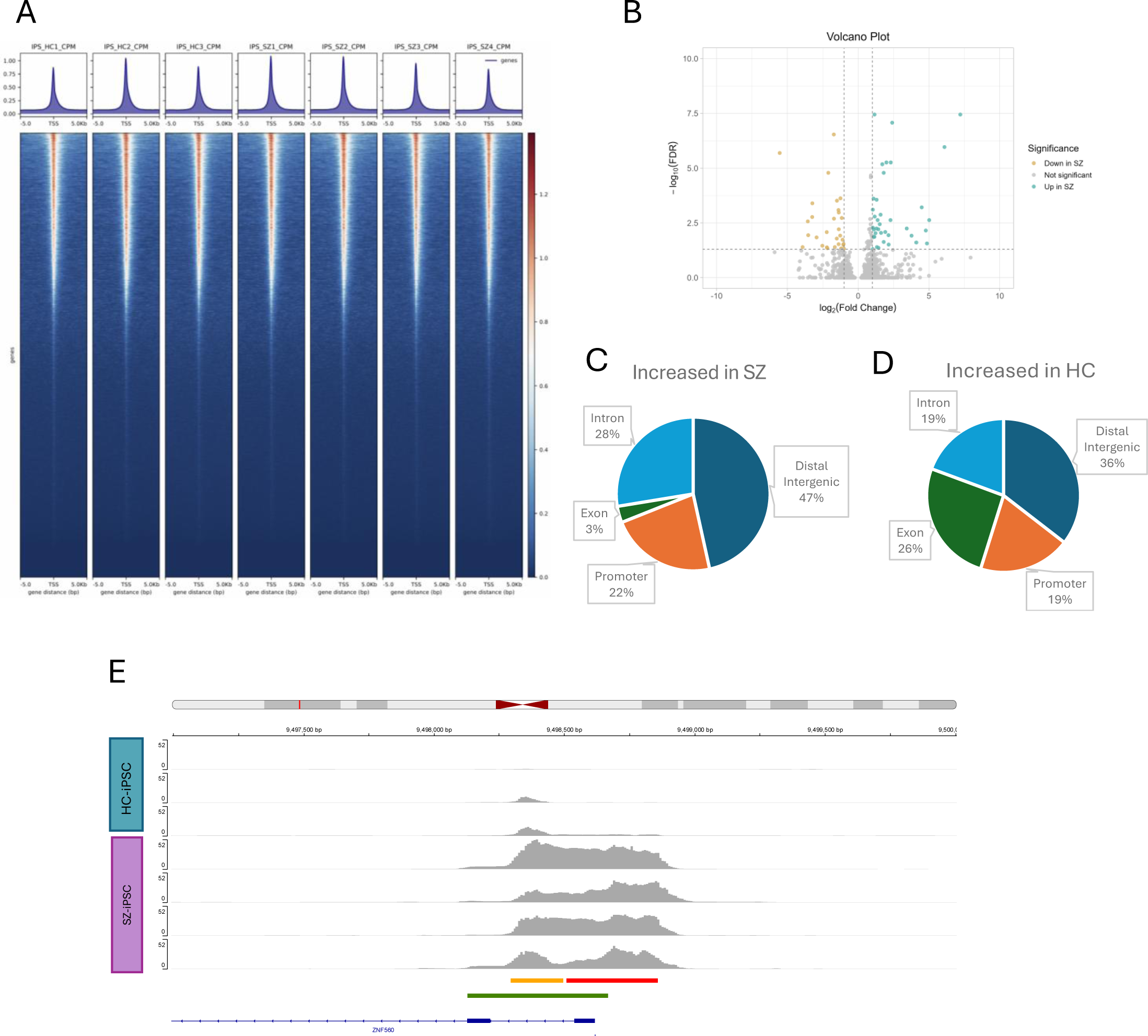
Differential chromatin accessibility in HC and SZ iPSC. **A.** Heatmaps of ATAC-seq profiles centered on transcription start sites (TSS ±5 kb) for individual iPSC samples (3 HC and 4 SZ) show globally comparable chromatin accessibility patterns across conditions. **B.** Volcano plot from differentially accessible regions (DARs) between SZ and HC iPSC. Regions with FDR < 0.05 and |log2 fold change| > 1 were considered differentially accessible. **C-D.** Genomic location of DARs showing the distribution of regions with increased (C) and decreased (D) accessibility in SZ. **E.** Genomic visualization of ATAC-seq signal tracks at the *ZNF5c0* promoter region in HC-iPSCs and SZ-iPSCs. Colored bars indicate the presence of Cis-Regulatory Elements (CREs) obtained from the ENCODE database: the red bar corresponds to promoter- like signatures and the yellow bar to proximal enhancer-like signatures. Green bar corresponds to a CpG island.

We further identified a significant increase in chromatin accessibility at a region spanning regulatory elements of the *ZNF560* gene promoter in SZ versus HC iPSC (Figure 2E). Using the “Find Individual Motif Occurrences” platform (FIMO), we then searched this region for conserved transcription factor motifs, identifying 75 significant associations (q< 0.01) for transcription factors (TFs) that could potentially bind to this region. Among this set of TFs, 53 exhibited transcript expression in iPSCs (Supplementary Figure 2A, Supplementary Table 7). Motifs for factors like KLF15, INSM1, and VDR were overrepresented within DARs associated with increased accessibility in SZ iPSC, whereas motifs for KLF9, SP3, and ZNF263 were identified within sequences associated with reduced accessibility in SZ iPSC (Supplementary Table 7 and 8). Overall, most of the TF motifs identified at the *ZNF560* locus localized to the gene promoter sequence of this gene (Supplementary Figure 2B).

### Epigenetic regulation of the ZNF5c0 promoter in HC and SZ iPSC

The *ZNF5c0* gene contains a CpG island encompassing both an enhancer and a promoter-like regulatory sequence [38, 39]. Because *de novo* DNA methylation is known to be acquired in the iPSC genome during reprogramming, we determined the DNA methylation status of the *ZNF5c0* locus (Figure 3A). This analysis showed a reduced CpG methylation percentage (1.6%) at the *ZNF560* promoter in fibroblasts, consistent with the active transcriptional status of the gene in these cells. This CpG methylation raised markedly to 91% in HC iPSC after reprogramming, consistent with *ZNF560* repression in these cells (Figure 3B-C). On the contrary, SZ iPSC exhibited a significantly lower CpG methylation rate (55%) at this region (Figure 3B-C), consistent with the persistence of active *ZNF560* transcription after reprogramming. Together, these results provide mechanistic support for the persistence of *ZNF560* gene expression in SZ-derived iPSC and suggest that an epigenetic-dependent mechanism mediating *ZNF5c0* gene repression is impaired during SZ fibroblast reprogramming to iPSC.

**Figure 3.**
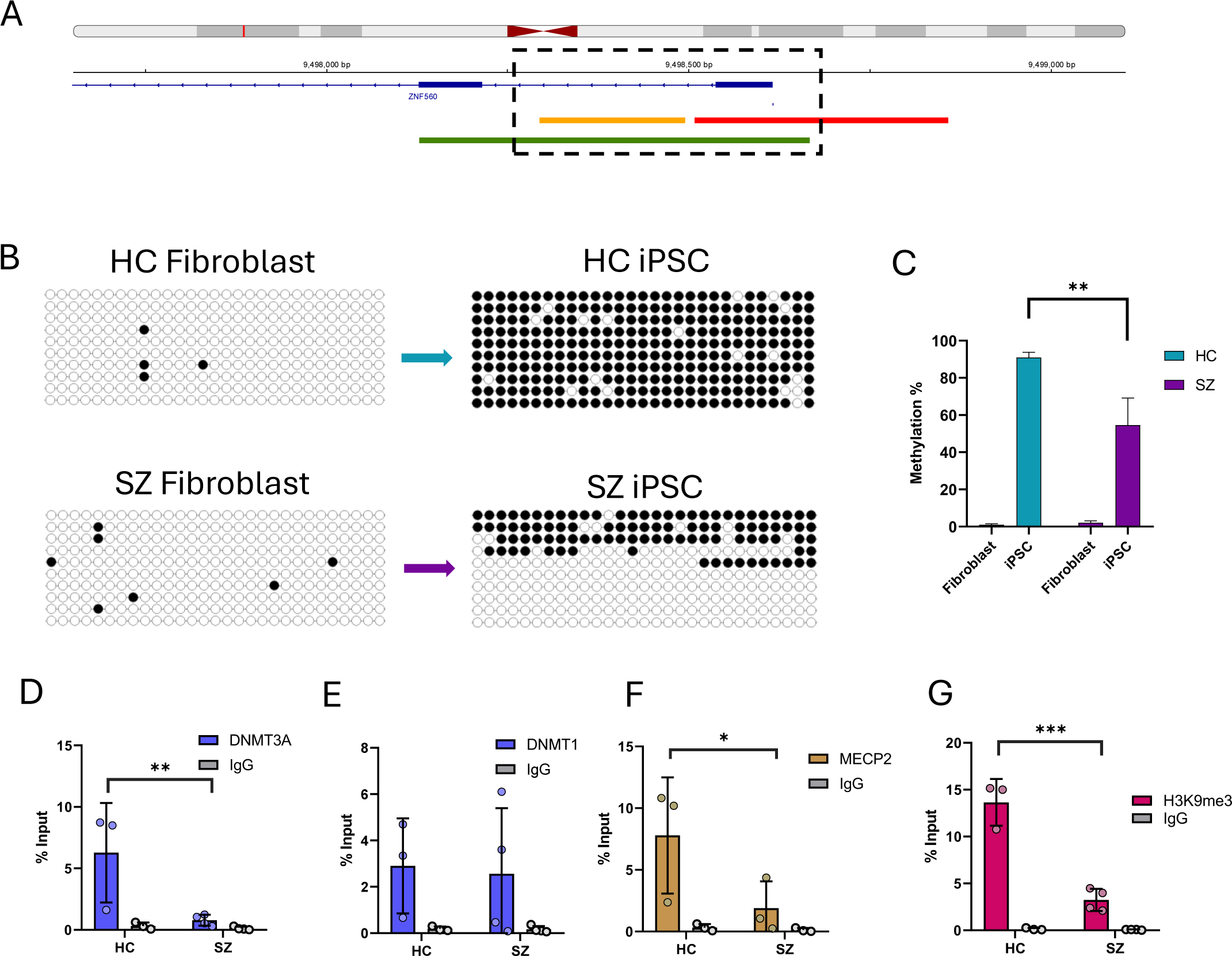
Impaired epigenetic silencing of ZNF560 in SZ iPSC. **A.** IGV visualization of ZNF560 gene promoter at chromosome 19. Red bar= promoter-like signatures; yellow bar= proximal enhancer-like signatures, green bar = CpG island. The DNA methylation levels at the CpG island that colocalizes with CREs (dashed area) were analyzed by bisulfite sequencing in Fibroblasts and iPSC. **B.** Bead diagram shows the presence (black circles) or absence (white circles) of methylation in cytosines of 29 CpG sites in representative HC and SZ iPSC samples and their isogenic fibroblasts. 10 clones were analyzed for each iPSC sample. **C.** The graph shows the methylation percentage for the whole sequence in each HC or SZ Fibroblast and iPSC (3 HC and 4 SZ). Data is shown as mean ± SD, with ** p<0.01 by 2-way ANOVA. **D-G.** The enrichment of DNMT3A (D), DNMT1 (E), MECP2 (F), and H3K9me3 (G) at this region was analyzed by ChIP–qPCR in iPSC (3 HC and 4 SZ). IgG was used as a negative control, and enrichment was calculated as a percentage of input. Data is shown as mean ± SD, with \**p* < 0.05; **p < 0.01; \*\*\**p* < 0.001 by 2-way ANOVA.

CpG methylation levels were assessed by bisulfite conversion followed by Sanger sequencing, with at least 10 clones sequenced per iPSC sample. As shown in Figure 3B, methylation percentages varied slightly among clones, potentially reflecting cell heterogeneity or monoallelic methylation. To further validate this analysis, we evaluated a non-related gene (*BGLAP*) known to be silent in non-osteoblastic cells and which exhibits limited, but consistent, CpG methylation at its silent promoter region [40]. Our results confirmed that the *BGLAP* promoter was fully methylated at these CpG sites in both HC and SZ iPSC genomes, further validating our experimental strategy to assess the DNA methylation status (Supplementary Figure 3A).

Because CpG methylation is mediated by a group of enzymes carrying methyltransferase activity, we next assessed the enrichment levels of DNA methyltransferases at the promoter region of *ZNF560* by ChIP-qPCR. It was found that at this region, the *de novo* methyltransferase *DNMT3A* showed significantly higher enrichment in HC compared to SZ iPSC, whereas no difference was detected for the maintenance methyltransferase *DNMT1* (Figure 3D-E). Because mRNA and protein levels of both *DNMTs* were comparable between HC and SZ iPSC (Supplementary Figure 3B-C), the reduced *DNMT3A* enrichment at the *ZNF560* promoter in SZ iPSC likely reflects reduced recruitment of this enzyme, rather than lower expression of the enzyme. In addition, no significant difference was detected in the expression (at the mRNA level) of the *TET* enzyme members *TET1*, *TET2*, and *TET3*, which catalyze the critical initial activities during the oxidative process that leads to CpG demethylation [41, 42] in mammals (Supplementary Figure 3C). We additionally assessed the presence of the methyl DNA-binding protein *MECP2*, which can recognize with high affinity methylated CpG dinucleotides and contribute to establishing compact chromatin conformations [43–45]. In agreement with the binding profile detected for *DNMT3A*, it was determined that *MECP2* exhibits higher enrichment at the *ZNF560* promoter in HC iPSC samples (Figure 3F). Importantly, *MECP2* transcript levels were comparable between HC and SZ iPSC samples (Supplementary Figure 3C), indicating that the differential enrichment profile between SZ and HC iPSC is not driven by differences in *MECP2* expression.

*MECP2* can mediate the recruitment of additional epigenetic modifiers that catalyze the deposition of histone repressive marks, including methylation at lysine 9 of histone 3 (H3K9me3) [46]. Therefore, we next measured the abundance of H3K9me3 at the *ZNF560* promoter in chromatin from both SZ and HC iPSC. H3K9me3 enrichment was elevated at the *ZNF5c0* gene promoter in HC iPSC (Figure 3G). This high enrichment was comparable to the H3K9me3 levels detected at other strongly repressed promoter region, the silent *SP7* gene (Supplementary Figure 3D). In contrast, H3K9me3 enrichment at the *ZNF560* gene promoter in SZ iPSC exhibited a significantly reduced level (Figure 3G), while H3K9me3 levels at the *SP7* gene promoter remained comparable between SZ and HC iPSC (Supplementary Figure 3D). Taken together, these results further suggest that an epigenetic repressive mechanism operates at the ZNF560 gene promoter during reprogramming from HC fibroblasts to iPSC. This epigenetically mediated repression is impaired when fibroblasts from SZ patients are subjected to reprogramming, resulting in persistent ZNF560 expression in SZ iPSC.

### Potential novel functions of ZNF5c0

ZNF560 belongs to the Krüppel-associated box (KRAB) domain-containing zinc finger proteins (KRAB-ZNF) family of transcriptional repressors. Members of this family are characterized by a KRAB domain at the N-terminal end and a C-terminal array of C2H2 zinc fingers [24]. KRAB-ZFNs bind to specific DNA regions through their zinc finger domains and recruit KAP1, which in turn mediates the subsequent interaction with several epigenetic modifiers to repress gene expression [23]. ChIP-seq studies show that these proteins primarily bind predominantly to transposable elements, silencing their expression during critical developmental windows and thereby regulating the expression of transcriptional networks [25, 26].

To identify additional putative DNA targets of *ZNF560*, we re-analyzed published ChIP-seq data from ZNF560 ectopically expressed in HEK293T cells (GEO accession: GSM6047482) [23]. Using the ChIPseeker package in R, we determined the genomic distribution of ZNF560 peaks, finding that 43% of its genomic binding sites are located at gene promoter regions (±2Kb from TSS) (Supplementary Figure 4A). GO analysis indicated that these bound target genes enriched biological processes that are mainly related to transcription regulation. Notably, several significantly associated terms were also associated with neural function, including nervous system development, neurogenesis, and neuron differentiation, among others (Supplementary Figure 4B). Additionally, cell compartment ontologies were also found enriched in terms related to neuronal compartments, and biological pathways ontologies included the “neuronal system” term (Supplementary Figure 4C-D). Because ZNF560 is predicted to act as a transcriptional repressor, we next asked whether its putative target genes overlap with genes downregulated in SZ NSC. We determined that 100 downregulated genes in SZ NSC (17 % of all downregulated genes) have the potential of binding *ZNF560* at their promoter region (Supplementary Figure 4E). GO analysis of these 100 genes showed association with neural-related terms, including: voltage-gated channel activity, glutamate receptor activity, trans-synaptic signaling, receptor localization to synapse, vesicle-mediated transport in synapse, synapse, and synaptic cleft (Supplementary Figure 4E).

Together, these results are consistent with a role of ZNF560 as a transcriptional regulator during early neurodevelopment. Moreover, the impaired control of ZNF560 repression in SZ iPSC argues in favor of potentially altered transcriptional control mechanisms affecting neuronal-related functions in SZ-derived iPSCs, a hypothesis that will require direct functional validation.

## Discussion

In this study, we have combined transcriptomic and epigenomic analyses of SZ-derived iPSC to identify early molecular alterations associated with the disease. While HC and SZ displayed highly similar global transcriptional profiles at the iPSC state, pronounced differences emerged upon neural lineage differentiation, supporting the view that schizophrenia-related molecular phenotypes are revealed during neurodevelopment rather than at the pluripotent stage [17, 47, 48].

SZ-NSC DEGs showed an enrichment of genes and pathways related to neuronal function. These findings are consistent with extensive evidence implicating synaptic dysfunction and neurotransmitter imbalance in schizophrenia pathophysiology [2, 3, 8, 9]. SynGO analysis further demonstrated that these alterations affect both pre- and postsynaptic compartments, including trans-synaptic signaling and postsynaptic density, closely mirroring signatures reported in *postmortem* schizophrenia cortex [2, 35, 49]. The partial overlap between NSC DEGs and *postmortem* DLPC genes reinforces the relevance of iPSC-based neurodevelopmental models in biomedical research, despite inherent differences in developmental stage and cellular context.

*ZNF560* emerged as a uniquely dysregulated gene in iPSC, showing persistent expression in SZ but effective silencing in HC iPSC following reprogramming. This pattern was robust across reprogramming methods, passages, and independent datasets, and demonstrated strong discriminatory power between HC and SZ iPSC, suggesting that *ZNF560* dysregulation reflects a stable disease-associated condition. Importantly, transcriptomic data of iPSC derived from patients with bipolar disorder (BD), a closely related disease to SZ with shared genetic alterations and overlapping symptoms, indicates that there is no significant difference in *ZNF5c0* expression between BD and HC samples, supporting the potential future use of *ZNF5c0* as a schizophrenia biomarker [50–52].

Epigenomic profiling of the promoter region of *ZNF560* revealed that its aberrant expression in SZ iPSC is associated with increased chromatin accessibility, reduced DNA methylation, and decreased DNMT3A and MECP2 recruitment, accompanied by decreased H3K9me3 enrichment. Our findings point to a locus-specific failure of epigenetic silencing rather than global alterations in methylation machinery and highlight *ZNF560* as a “reprogramming-resistant” gene in schizophrenia. Interestingly, a recent study in Alzheimer’s disease (AD) also described epigenomic alterations in AD-iPSC that were associated with functional imbalances [53]. Together, these data challenge our long-standing knowledge of epigenetic reprogramming in iPSC and proposes the existence of potential disease-associated epigenetic impairments.

Among epigenetic regulators, KRAB-ZNFs represent the largest family of transcriptional repressors in vertebrates, with 378 members encoded in the human genome. They bind to specific DNA motifs and recruit KRAB-associated protein 1 (KAP1), promoting epigenetic silencing through chromatin-modifying complexes [23, 24]. KRAB-ZNFs are highly expressed during early development and have been implicated in the regulation of transposable elements and gene networks controlling cell fate decisions [25, 26]. Notably, several KRAB-ZNFs are expressed during brain development and have been shown related to processes like neurogenesis, gliogenesis, NSC proliferation, synaptic transmission, and neuronal death, among others [24, 54]. However, the specific mechanisms underlying these regulatory functions have not been sufficiently uncovered. Therefore, the impact of an altered expression of these factors in nervous system disorders remains undetermined. Our analysis of published ZNF560 ChIP-seq data revealed prominent binding at gene promoters, including genes implicated in neurodevelopment and synaptic function. Notably, a substantial fraction of genes downregulated in SZ NSC harbor *ZNF560* binding sites and are associated with GO terms including synaptic signaling and neurotransmission-related processes, hence suggesting that persistent *ZNF5c0* expression may contribute to impaired neuronal gene regulation during differentiation. However, because the ZNF560 ChIP-seq dataset was generated in HEK293 cells rather than in neural cells, these findings should be validated in a neurodevelopmental context; nevertheless, they provide a valuable and biologically coherent framework to support our conclusions.

In summary, our results identify *ZNF560* as a dysregulated gene in schizophrenia-derived iPSC, linked to defective epigenetic silencing during fibroblast reprogramming and associated with downstream transcriptional alterations in NSC. These results highlight a previously unrecognized role for a KRAB-ZNF protein in schizophrenia-related neurodevelopmental pathways and support the utility of iPSC-based models for uncovering stable, disease-associated epigenetic mechanisms.

## Acknowledgments

Funding supporting this work: Millennium Nucleus of Neuroepigenetics and Plasticity NCN2023_032 (MM), ANID-FONDECYT 1251450 (MM), ANID-FONDECYT 1221522 (VP), ANID-FONDECYT POSTDOCTORADO 3230411 (BSC), and ANID-FONDECYT INICIACION 11260455 (BSC).

## References

1. Owen MJ, Legge SE, Rees E, Walters JTR, O’Donovan MC. Genomic findings in schizophrenia and their implications. Mol Psychiatry. 2023;28:3638–3647.

2. Trubetskoy V, Pardiñas AF, Qi T, Panagiotaropoulou G, Awasthi S, Bigdeli TB, et al. Mapping genomic loci implicates genes and synaptic biology in schizophrenia. Nature. 2022;604:502 – 508.

3. Hall J, Bray NJ. Schizophrenia Genomics: Convergence on Synaptic Development, Adult Synaptic Plasticity, or Both? Biol Psychiatry. 2022;91:709–717.

4. Stilo SA, Di Forti M, Murray RM. Environmental risk factors for schizophrenia: implications for prevention. Neuropsychiatry. 2011;1:457–466.

5. Schmidt-Kastner R, van Os J, Esquivel G, Steinbusch HWM, Rutten BPF. An environmental analysis of genes associated with schizophrenia: hypoxia and vascular factors as interacting elements in the neurodevelopmental model. Mol Psychiatry. 2012;17:1194–1205.

6. Yang H, Sun W, Li J, Zhang X. Epigenetics factors in schizophrenia: future directions for etiologic and therapeutic study approaches. Ann Gen Psychiatry. 2025;24:21.

7. Richetto J, Meyer U. Epigenetic Modifications in Schizophrenia and Related Disorders: Molecular Scars of Environmental Exposures and Source of Phenotypic Variability. Biol Psychiatry. 2021;89:215–226.

8. Faludi G, Mirnics K. Synaptic changes in the brain of subjects with schizophrenia. International Journal of Developmental Neuroscience. 2011;29:305–309.

9. Howes OD, Onwordi EC. The synaptic hypothesis of schizophrenia version III: a master mechanism. Mol Psychiatry. 2023;28:1843–1856.

10. De Los Angeles A, Fernando MB, Hall NAL, Brennand KJ, Harrison PJ, Maher BJ, et al. Induced Pluripotent Stem Cells in Psychiatry: An Overview and Critical Perspective. Biol Psychiatry. 2021;90:362–372.

11. Moslem M, Olive J, Falk A. Stem cell models of schizophrenia, what have we learned and what is the potential? Schizophr Res. 2019;210:3–12.

12. Karagiannis P, Takahashi K, Saito M, Yoshida Y, Okita K, Watanabe A, et al. Induced pluripotent stem cells and their use in human models of disease and development. Physiol Rev. 2019;99:79–114.

13. Stachowiak EK, Benson CA, Narla ST, Dimitri A, Chuye LEB, Dhiman S, et al. Cerebral organoids reveal early cortical maldevelopment in schizophrenia—computational anatomy and genomics, role of FGFR1. Transl Psychiatry. 2017;7:6.

14. Nascimento JM, Saia-Cereda VM, Zuccoli GS, Reis-de-Oliveira G, Carregari VC, Smith BJ, et al. Proteomic signatures of schizophrenia-sourced iPSC-derived neural cells and brain organoids are similar to patients’ postmortem brains. Cell Biosci. 2022;12:1–16.

15. Da Silveira Paulsen B, De Moraes Maciel R, Galina A, Da Silveira MS, Souza CDS, Drummond H, et al. Altered Oxygen Metabolism Associated to Neurogenesis of Induced Pluripotent Stem Cells Derived from a Schizophrenic Patient. Cell Transplant. 2012;21:1547–1559.

16. Brennand K, Savas JN, Kim Y, Tran N, Simone A, Hashimoto-Torii K, et al. Phenotypic differences in hiPSC NPCs derived from patients with schizophrenia. Mol Psychiatry. 2015;20:361–368.

17. Brennand KJ, Simone A, Jou J, Gelboin-Burkhart C, Tran N, Sangar S, et al. Modelling schizophrenia using human induced pluripotent stem cells. Nature. 2011;473:221.

18. Casas BS, Vitória G, do Costa MN, Madeiro da Costa R, Trindade P, Maciel R, et al. hiPSC-derived neural stem cells from patients with schizophrenia induce an impaired angiogenesis. Transl Psychiatry. 2018;8:48.

19. Puvogel S, Blanchard K, Casas BS, Miller RL, Garrido-Jara D, Arizabalos S, et al. Altered resting-state functional connectivity in hiPSCs-derived neuronal networks from schizophrenia patients. Front Cell Dev Biol. 2022;10.

20. Trindade P, Nascimento JM, Casas BS, Monteverde T, Gasparotto J, Ribeiro CT, et al. Induced pluripotent stem cell-derived astrocytes from patients with schizophrenia exhibit an inflammatory phenotype that affects vascularization. Mol Psychiatry. 2023;28:871–882.

21. Lee S-A, Huang K-C. Epigenetic profiling of human brain differential DNA methylation networks in schizophrenia. BMC Med Genomics. 2016;9:68.

22. Guidotti A, Auta J, Davis JM, Dong E, Gavin DP, Grayson DR, et al. Toward the Identification of Peripheral Epigenetic Biomarkers of Schizophrenia. J Neurogenet. 2014;28:41–52.

23. de Tribolet-Hardy J, Thorball CW, Forey R, Planet E, Duc J, Coudray A, et al. Genetic features and genomic targets of human KRAB-zinc finger proteins. Genome Res. 2023;33:1409–1423.

24. Ecco G, Imbeault M, Trono D. KRAB zinc finger proteins. Development. 2017;144:2719– 2729.

25. Farmiloe G, Lodewijk GA, Robben SF, van Bree EJ, Jacobs FMJ. Widespread correlation of KRAB zinc finger protein binding with brain-developmental gene expression patterns. Philosophical Transactions of the Royal Society B: Biological Sciences. 2020;375:20190333.

26. Chen Y-C, Maupas A, Nowick K. Regulatory networks of KRAB zinc finger genes and transposable elements changed during human brain evolution and disease. Elife. 2025.

27. Bock C, Reither S, Mikeska T, Paulsen M, Walter J, Lengauer T. BiQ Analyzer: visualization and quality control for DNA methylation data from bisulfite sequencing. Bioinformatics. 2005;21:4067–4068.

28. Soutoglou E, Talianidis I. Coordination of PIC Assembly and Chromatin Remodeling During Differentiation-Induced Gene Activation. Science (1979). 2002;295:1901–1904.

29. Pola-Véliz V, Arredondo SB, Arancibia Y, Ahumada J, Estay S, Vidal N, et al. Viral-mediated fluorescent labelling of activated hippocampal memory engrams to study epigenetic dynamics associated with gene expression. Methods. 2026;247:85–94.

30. Berretta S. Extracellular matrix abnormalities in schizophrenia. Neuropharmacology. 2012;62:1584–1597.

31. Pantazopoulos H, Katsel P, Haroutunian V, Chelini G, Klengel T, Berretta S. Molecular signature of extracellular matrix pathology in schizophrenia. European Journal of Neuroscience. 2021;53:3960–3987.

32. Tsimberidou A-M, Skliris A, Valentine A, Shaw J, Hering U, Vo HH, et al. AKT inhibition in the central nervous system induces signaling defects resulting in psychiatric symptomatology. Cell Biosci. 2022;12:56.

33. Enriquez-Barreto L, Morales M. The PI3K signaling pathway as a pharmacological target in Autism related disorders and Schizophrenia. Mol Cell Ther. 2016;4:2.

34. Koopmans F, van Nierop P, Andres-Alonso M, Byrnes A, Cijsouw T, Coba MP, et al. SynGO: An Evidence-Based, Expert-Curated Knowledge Base for the Synapse. Neuron. 2019;103:217–234.e4.

35. Fromer M, Roussos P, Sieberts SK, Johnson JS, Kavanagh DH, Perumal TM, et al. Gene expression elucidates functional impact of polygenic risk for schizophrenia. Nat Neurosci. 2016;19:1442–1453.

36. Klemm SL, Shipony Z, Greenleaf WJ. Chromatin accessibility and the regulatory epigenome. Nat Rev Genet. 2019;20:207–220.

37. Hsieh J, Gage FH. Chromatin remodeling in neural development and plasticity. Curr Opin Cell Biol. 2005;17:664–671.

38. Abascal F, Acosta R, Addleman NJ, Adrian J, Afzal V, Ai R, et al. Expanded encyclopaedias of DNA elements in the human and mouse genomes. Nature. 2020;583:699–710.

39. Moore JE, Pratt HE, Fan K, Phalke N, Fisher J, Elhajjajy SI, et al. An expanded registry of candidate cis-regulatory elements. Nature. 2026. 7 January 2026. 10.1038/s41586-025-09909-9.

40. Kang M, Kim H, Jung Y, Kim Y, Hong S, Kim M, et al. Transitional CpG methylation between promoters and retroelements of tissue-specific genes during human mesenchymal cell differentiation. J Cell Biochem. 2007;102:224–239.

41. Zhang X, Zhang Y, Wang C, Wang X. TET (Ten-eleven translocation) family proteins: structure, biological functions and applications. Signal Transduct Target Ther. 2023;8:297.

42. Rasmussen KD, Helin K. Role of TET enzymes in DNA methylation, development, and cancer. Genes Dev. 2016;30:733–750.

43. Boxer LD, Renthal W, Greben AW, Whitwam T, Silberfeld A, Stroud H, et al. MeCP2 Represses the Rate of Transcriptional Initiation of Highly Methylated Long Genes. Mol Cell. 2020;77:294–309.e9.

44. Nan X, Campoy FJ, Bird A. MeCP2 Is a Transcriptional Repressor with Abundant Binding Sites in Genomic Chromatin. Cell. 1997;88:471–481.

45. Pantier R, Brown M, Han S, Paton K, Meek S, Montavon T, et al. MeCP2 binds to methylated DNA independently of phase separation and heterochromatin organisation. Nat Commun. 2024;15:3880.

46. Fuks F, Hurd PJ, Wolf D, Nan X, Bird AP, Kouzarides T. The Methyl-CpG-binding Protein MeCP2 Links DNA Methylation to Histone Methylation. Journal of Biological Chemistry. 2003;278:4035–4040.

47. Hoffman GE, Hartley BJ, Flaherty E, Ladran I, Gochman P, Ruderfer DM, et al. Transcriptional signatures of schizophrenia in hiPSC-derived NPCs and neurons are concordant with post-mortem adult brains. Nat Commun. 2017;8.

48. Hoffman GE, Schrode N, Flaherty E, Brennand KJ. New considerations for hiPSC-based models of neuropsychiatric disorders. Mol Psychiatry. 2019;24:49–66.

49. Ruzicka WB, Mohammadi S, Fullard JF, Davila-Velderrain J, Subburaju S, Tso DR, et al. Single-cell multi-cohort dissection of the schizophrenia transcriptome. Science (1979). 2024;384.

50. Madison JM, Zhou F, Nigam A, Hussain A, Barker DD, Nehme R, et al. Characterization of bipolar disorder patient-specific induced pluripotent stem cells from a family reveals neurodevelopmental and mRNA expression abnormalities. Mol Psychiatry. 2015;20:703– 717.

51. Chen HM, DeLong CJ, Bame M, Rajapakse I, Herron TJ, McInnis MG, et al. Transcripts involved in calcium signaling and telencephalic neuronal fate are altered in induced pluripotent stem cells from bipolar disorder patients. Transl Psychiatry. 2014;4:e375–e375.

52. Shao L, Vawter MP. Shared Gene Expression Alterations in Schizophrenia and Bipolar Disorder. Biol Psychiatry. 2008;64:89–97.

53. Katbe A, Hanna R, Flamier A, Serhani D, Hamam R, Barabino A, et al. Epigenomic alterations and neural development anomalies in induced pluripotent stem cells from sporadic Alzheimer’s disease. Development. 2026;153.

54. Al-Naama N, Mackeh R, Kino T. C2H2-Type Zinc Finger Proteins in Brain Development, Neurodevelopmental, and Other Neuropsychiatric Disorders: Systematic Literature-Based Analysis. Front Neurol. 2020;11.

55. Casas BS, Vitória G, Prieto CP, Casas M, Chacón C, Uhrig M, et al. Schizophrenia-derived hiPSC brain microvascular endothelial-like cells show impairments in angiogenesis and blood–brain barrier function. Mol Psychiatry. 2022;27:3708–3718.

