## Supplementary Information for "Impaired ZNF560 repression during induced pluripotent stem cell reprogramming indicates early epigenetic alterations in Schizophrenia"

1. Supplemental Methods
2. Supplementary Table 1
3. Supplementary Table 2
4. Supplementary Table 3
5. Supplementary Table 5
6. Supplementary Table 7
7. Supplementary Table 8
8. Supplmentary Table 9
9. Supplementary Figure 1
10. Supplementary Figure 2
11. Supplementary Figure 3
12. Supplementary Figure 4

Supplemental Methods

*Cell lines*

Detailed information on patients and the source of cells is provided in Supplementary Table 1. We selected SZP iPSC that came from patients with known familial risk of SZ, ensuring a high genetic component. For expansion, iPSC were seeded in Matrigel-coated plates and cultured in mTESR1 medium with daily medium changes (StemCells Technologies Cat N° 85850, Canada), following the manufacturer's instructions.

Isogenic fibroblasts (source of iPSC reprogramming) were cultured in DMEM high glucose supplemented with 10% Fetal Bovine Serum as described in [1]. Medium was changed every other day.

Four iPSC lines (two from control subjects and two from schizophrenia patients) were differentiated into NSC following the protocol described in [1]. Briefly, iPSC cells were cultured in Pluripotent Stem Cells (PSC) Neural Induction Medium (Thermo Fisher Scientific, Carlsbad, CA, USA), changing medium every 48 hours for 7 days. Cells were split and then cultured in Neural Induction Medium (NEM, Advanced DMEM/F12 and Neurobasal medium (1:1) with Neural Induction Supplement; Thermo Fisher Scientific, Carlsbad, CA, USA). Cultures were used between passages 14 and 18.

*RNA extraction and RT-qPCR*

Cells were homogenized in Trizol, and then total RNA was extracted using Direct-zol RNA MiniPrep (Zymo Research Cat No R2052, Irvine, CA).

1 µg of total RNA was retrotranscribed using M-MuLV Reverse Transcriptase (NewEngland Biolabs, M0253S, Ontario, Canada). Relative expression was assessed by qPCR using

SybrGreen II mix (Agilent Technologies Thermocycler, Santa Clara, CA, USA) and specific primers listed in Supplementary Table 2. Data was analyzed by calculating the expression fold change via $2^{-\Delta\Delta Ct}$, and gene expression was normalized to that of three reference genes (*GAPDH, B2M, and C1orf43*).

*RNA-seq*

1 µg of total RNA was sent to Quick Biology (https://www.quickbiology.com/; Monrovia, CA, USA) for a standard RNA sequencing service, PolyA, 30M reads per sample, paired-end.

Raw sequencing reads were assessed for quality using FastQC (v0.11.9) and screened for contamination with FastQ Screen. Adapter sequences and low-quality bases were trimmed using Trim Galore (v0.6.7). Cleaned reads were aligned to the human reference genome (hg38, ENCODE v3) using STAR (v2.7.10a) with default parameters for splice-aware alignment. Gene-level read counts were quantified using featureCounts (v2.0.1) with GENCODE annotation. Differential expression analysis between schizophrenia (SZ) and healthy control (HC) iPSC-derived samples was performed using DESeq2 (v1.34.0) in R. Genes with an adjusted p-value $< 0.05$ and $|\log 2 \text{ fold change}| > 1$ were considered differentially expressed. A variance-stabilizing transformation (VST) was applied for visualization and downstream analyses.

Publicly available datasets used in this article are listed in (Supplementary Table 3).

*ATAC-seq*

$6*10^5$ cells were cryopreserved in mTeSR1 with 5% DMSO and sent to Quick Biology for a standard ATAC-seq service, 50M reads, paired-end.

ATAC-seq data quality was evaluated using FastQC and FastQ Screen. Nextera adapter sequences and low-quality bases were removed using Trim Galore. Reads were aligned to the human reference genome (hg38, ENCODE v3) using Bowtie2 (v2.4.5) with parameters optimized for ATAC-seq (-X 2000 --very-sensitive --no-discordant --no-mixed). Aligned reads were sorted and indexed using SAMtools (v1.15). PCR duplicates were marked using Picard MarkDuplicates and subsequently removed. Reads mapping to mitochondrial DNA and ENCODE blacklisted regions were filtered out. Peak calling was performed using MACS2 (v2.2.7) with parameters -f BAMPE --nomodel --keep-dup all -q 0.1 --min-length 100 for both narrow and broad peak detection. Signal tracks were generated and normalized using MACS2 bdgcmp and converted to bigWig format for visualization. Differentially accessible regions obtained with DiffBind/DESeq2 (ATAC-seq; HC vs SZ contrast).

*Integration of GWAS and ATAC-seq data*

To identify differentially accessible chromatin regions overlapping schizophrenia risk loci, we intersected our ATAC-seq differential peaks with SNP positions from the Psychiatric Genomics Consortium (PGC3) schizophrenia GWAS. For each of the 287 genome-wide significant loci, we used the top associated SNP position (top-pos) as the reference coordinate. Differential ATAC-seq peaks (identified by DiffBind/DESeq2, SZ vs HC) were intersected with these SNP positions using coordinate overlap in R. A peak was considered overlapping if the SNP position fell within the peak boundaries (peak_start ≤ SNP_position ≤ peak_end). Peaks were annotated by direction of change (increased or decreased accessibility in schizophrenia).

*Motif enrichment analysis*

To identify de novo motifs with enriched signal in ATAC-seq peaks, motif enrichment analysis was performed using HOMER's findMotifsGenome.pl, comparing HC-up and SZ-up target sets against their respective backgrounds with hg38 as reference genome. Analysis was performed on 200 bp windows centered on each peak, and de novo motif discovery was restricted to lengths of 8, 10, 12, and 15 bp. HOMER reported an enrichment P-value for each motif (target vs background comparison) and putative transcription factor assignments based on similarity to integrated motif databases.

*FIMO analysis*

To generate a unified reference for sequence scanning, a motif library in MEME format (all_PWMs_hg38.meme) was constructed by concatenating human PWMs from HOCOMOCO v11 (core/full, as available) and JASPAR databases. FIMO (MEME-suite) was used to scan specific regions of the hg38 genome on both strands, evaluating motif matches by calculating per-occurrence p-values and their correction for multiple testing. Results were filtered using an FDR <0.01 threshold. The analyzed sequences corresponded to the chr19:9497998–9498917 interval on chromosome 19 of hg38.

*Methylation sequencing*

Genomic DNA was extracted using Quick-DNA MiniPrep (Zymo Research Cat. No D3024, Irvine, CA). To identify methylation of 5mC in DNA samples, 500 ng of DNA was bisulfite-converted using EZ DNA Methylation-Gold Kit (Zymo Research Cat. No D5005, Irvine, CA).

Regions of interest were amplified by PCR using specific converted primers (Supplementary Table 2). The PCR product was purified and cloned with a commercial TOPO TA Cloning Kit (Invitrogen Cat No: 450641, Carlsbad, CA). Sequences were obtained by Sanger method using M13 primers. Sequences were analyzed using BiQ Analyzer [2].

*Chomatin immunoprecipitation (ChIP)-PCR*

ChIP was performed as described previously [3, 4], with minor modifications. Cells were crosslinked with 1% formaldehyde, quenched with glycine, and nuclei were isolated by hypotonic lysis (50 mM HEPES pH 7.8, 3 mM $MgCl_2$, 20 mM KCl, 0.1% NP-40, 1x protease inhibitor cocktail) followed by mechanical disruption using a Dounce homogenizer with a tight pestle (Sigma-Aldrich, D8938). Chromatin was sonicated to 200–500 bp fragments in sonication buffer (50 mM HEPES pH 7.8, 140 mM NaCl, 1 mM EDTA, 1% Triton X-100, 0.1% SDS, 0.1% sodium deoxycholate, 1x protease inhibitors) using a Bioruptor and pre-cleared with isotype-matched IgG (rabbit IgG Millipore, 12-370 or mouse IgG Sigma, 12-371) and Dynabeads (sheep anti-rabbit Invitrogen 11203D or sheep anti-mouse Invitrogen 11202D). Chromatin from $1\times10^6$ nuclei was immunoprecipitated overnight at 4°C with antibodies against DNMT1 (Abcam, ab13537) , DNMT3A (Abcam, ab2850), MECP2 (Abcam, ab2828), H3K9me3 (Abcam, ab8898), followed by capture with Dynabeads. Immunocomplexes were washed sequentially, eluted, and reverse crosslinked. DNA was purified using ChIP DNA Clean & Concentrator columns (Zymo Research, D5205), and enrichment was quantified by qPCR using specific primers (Supplementary Table 2).

*Western blot*

For nuclear protein extraction cells were lysed in ice-cold Buffer A (10 mM Hepes pH 7.9, 5 mM $MgCl_2$, 0.25M Sucrose) supplemented with 0.1% NP-40 and protease/phosphatase inhibitors to release the cytosolic fraction. After centrifugation, intact nuclei were resuspended in high-salt Buffer B (25 mM Hepes pH 7.9, 20%, glycerol, 1.5 mM $MgCl_2$, 0.1 mM EDTA, NaCl to 700mM) containing protease/phosphatase inhibitors and incubated on ice, followed by brief sonication to solubilize chromatin-bound proteins. Nuclear extracts were clarified by high-speed centrifugation, and the supernatant containing nuclear and chromatin-associated proteins was collected and stored at −80 °C until use.

Western blotting was performed as described before [1, 5]. Membranes were incubated with DNMT1 (Abcam, ab13537) or DNMT3A (Abcam, ab2850) antibodies overnight at 4 °C. The monoclonal mouse anti-PCNA (13-39000, Invitrogen) antibody was used as a loading control. Membranes were washed and incubated (2 h, 22 °C) with horseradish peroxidase-conjugated goat anti-mouse secondary antibodies. Bands were visualized using enhanced chemiluminescence (ECL; Amersham Biosciences, Little Chalfont, UK), and quantified by densitometry using ImageJ (NIH, USA).

*Data availability*

RNA-seq datasets have been deposited in Gene Expression Omnibus GSE325013 and ATAC-seq in GSE325014.

| | | | | | | Use in this study | | | | |
|---|---|---|---|---|---|---|---|---|---|---|
| **ID** | **Diagnosis** | **Age** | **Sex** | **Reprogramming Technique** | **Code and references** | **RNA -seq** | **RT-qPCR** | **ATAC -seq** | **BS-seq & ChIP** | **Diff. into NSC** |
| HC1 | Healthy control | F | 36 | Retroviral vectors (OCT4, SOX2, KLF4, CMYC ) | Coriell Institute N°GM23279 | x | x | x | x | |
| HC2 | Healthy control | M | 37 | CytoTune®-iPS 2.0 Kit (Thermo Fisher) | CF1 [1, 5–8] | x | x | x | x | x |
| HC3 | Healthy control | M | 31 | CytoTune®-iPS 2.0 Kit (Thermo Fisher) | CF2 [1, 5–7] | x | x | x | x | x |
| HC4 | Healthy control | M | 64 | Lentiviral vector (OCT4, SOX2, KLF4, CMYC, LIN28) | Control 37L25 [9] | | x | | | |
| HC5 | Healthy control | M | 64 | Lentiviral vector (OCT4, SOX2, KLF4, CMYC, LIN28) | Control [10] | | x | | | |
| HC6 | Healthy control | F | neon ate | Episomal vector (Oct4, Sox2, Klf4, Myc, Nanog, Lin28, and SV40 T) | Gibco™ episomal hiPSC line A18945 | | x | | | |
| SZ1 | Schizophrenia | F | 27 | Lentiviral vector (OCT4, SOX2, KLF4, CMYC, LIN28) | Coriell Institute N°GM23761 | x | x | x | x | x |
| SZ2 | Schizophrenia | M | 26 | Lentiviral vector (OCT4, SOX2, KLF4, CMYC, LIN28) | Coriell Institute N°GM23760 | x | x | x | x | x |
| SZ3 | Schizophrenia | M | 23 | Lentiviral vector (OCT4, SOX2, KLF4, CMYC, LIN28) | Coriell Institute N°GM23762 | x | x | x | x | |
| SZ4 | Schizophrenia | M | 45 | CytoTune®-iPS 2.0 Kit (Thermo Fisher) | EZQ3 [11] | x | x | x | x | |
| SZ5 | Schizophrenia | M | 44 | CytoTune®-iPS 2.0 Kit (Thermo Fisher) | EZQ4 [11] | | x | | | |
| SZ6 | Schizophrenia | M | 44 | CytoTune®-iPS 2.0 Kit (Thermo Fisher) | EZQ9 [11] | | x | | | |

**Supplementary Table 1.** iPSC lines used in this study. ID: identification in the manuscript.


1. Casas BS, Vitória G, do Costa MN, Madeiro da Costa R, Trindade P, Maciel R, et al. hiPSC-derived neural stem cells from patients with schizophrenia induce an impaired angiogenesis. Transl Psychiatry. 2018;8:48.
2. Bock C, Reither S, Mikeska T, Paulsen M, Walter J, Lengauer T. BiQ Analyzer: visualization and quality control for DNA methylation data from bisulfite sequencing. Bioinformatics. 2005;21:4067–4068.
3. Soutoglou E, Talianidis I. Coordination of PIC Assembly and Chromatin Remodeling During Differentiation-Induced Gene Activation. Science (1979). 2002;295:1901–1904.

4. Pola-Véliz V, Arredondo SB, Arancibia Y, Ahumada J, Estay S, Vidal N, et al. Viral-mediated fluorescent labelling of activated hippocampal memory engrams to study epigenetic dynamics associated with gene expression. Methods. 2026;247:85–94.
5. Casas BS, Vitória G, Prieto CP, Casas M, Chacón C, Uhrig M, et al. Schizophrenia-derived hiPSC brain microvascular endothelial-like cells show impairments in angiogenesis and blood–brain barrier function. Mol Psychiatry. 2022;27:3708–3718.
6. Ledur PF, Karmirian K, Pedrosa C da SG, Souza LRQ, Assis-de-Lemos G, Martins TM, et al. Zika virus infection leads to mitochondrial failure, oxidative stress and DNA damage in human iPSC-derived astrocytes. Sci Rep. 2020;10:1218.
7. Assis-de-Lemos G, Monteiro J, Oliveira-Valença VM, Melo GA, Reis RA de M, Rehen SK, et al. Dopamine signaling impairs ROS modulation by mitochondrial hexokinase in human neural progenitor cells. Biosci Rep. 2021;41.
8. Puvogel S, Blanchard K, Casas BS, Miller RL, Garrido-Jara D, Arizabalos S, et al. Altered resting-state functional connectivity in hiPSCs-derived neuronal networks from schizophrenia patients. Front Cell Dev Biol. 2022;10.
9. Zhang Z, Almeida S, Lu Y, Nishimura AL, Peng L, Sun D, et al. Downregulation of MicroRNA-9 in iPSC-Derived Neurons of FTD/ALS Patients with TDP-43 Mutations. PLoS One. 2013;8:e76055-.
10. Almeida S, Zhang Z, Coppola G, Mao W, Futai K, Karydas A, et al. Induced Pluripotent Stem Cell Models of Progranulin-Deficient Frontotemporal Dementia Uncover Specific Reversible Neuronal Defects. Cell Rep. 2012;2:789–798.
11. Sochacki J, Devalle S, Reis M, Fontenelle LF, Rehen S. Generation of urine iPS cell line from a patient with obsessive-compulsive disorder using a non-integrative method. Stem Cell Res. 2016;17:107–110.

| Gene | Objective | FW | RV |
|---|---|---|---|
| ZNF560 | RT-qPCR | GGGATACAGTTGGCAAGAA | AGAAAGGAGGTCTTTTGCAT |
| DNMT1 | RT-qPCR | GTCCTTGGAGAACGGTGCT | CTTGAACGCTTAGCCTCTCCA |
| DNMT3A | RT-qPCR | CAGGCCGCATTGTGTCTTG | TGCTTGTTGTACGTGGCCT |
| MECP2 | RT-qPCR | CTTCTGCCTCCCCCAAACA | AAGGCTTTTCCCTGGGGATTG |
| TET1 | RT-qPCR | TGTTGTGCCTCTGGAGGTT | TGTTCTTCCCCATGACCACA |
| TET2 | RT-qPCR | ACTCGGTCAATTTCCCAGT | CGCGTCTCTATCTCCCTTT |
| TET3 | RT-qPCR | CTGAAGGGTGGATTGTCCC | GGTAGGAGTACACGCTGTTC |
| GAPDH | RT-qPCR | CAAGAAGGTGGTGAAGCAGGC | CCACCACCCTGTTGCTGTAG |
| B2M | RT-qPCR | GCTATCCAGCGTACTCCAAA | GAAAGACCAGTCCTTGCTGA |
| C1orf43 | RT-qPCR | ACGCCTTTCAAGGGTGTACG | CAAAGACCCCTGTCCCATAGC |
| ZNF560 | BS-seq | TTGTTTTGGAGATTTATTTTTT | AATCCTAAAAAATTCCATCC |
| BGLAP | BS-seq | AGGGTAGGGTTTGAGTCGT | GGGGGTATTGCGAGGTAT |
| ZNF560 | ChIP | TCGATTACCCTGGGTCGATT | TACTTTGGTGAGGCTCGTGT |
| GAPDH | ChIP | TGCTCCTCCTGTTCGGACAGT | CTAGCCTCCCGGGTTTCTCT |
| SP7 | ChIP | TTCATGATAGTGGCTGTACCCATG | ACTGGAACGCACTGACATACA |

**Supplementary Table 2.** Primers used in this study.

| Repository | Cell/Dataset identification | Diagnostic |
|---|---|---|
| this study | SZ1 | Schizophrenia* |
| this study | SZ2 | Schizophrenia* |
| this study | SZ3 | Schizophrenia* |
| this study | SZ4 | Schizophrenia* |
| SRA | SRX13198487 | Schizophrenia |
| SRA | SRX13198355 | Schizophrenia |
| SRA | SRX13198518 | Schizophrenia |
| this study | HC1 | Healthy Control* |
| this study | HC2 | Healthy Control* |
| this study | HC3 | Healthy Control* |
| this study | HC4 | Healthy Control* |
| this study | HC5 | Healthy Control* |
| this study | HC6 | Healthy Control* |
| SRA | SRX13198513 | Healthy Control |
| SRA | SRX13198294 | Healthy Control |
| SRA | SRX13198299 | Healthy Control |
| GEO | GSM2072610 | Healthy Control |
| GEO | GSM2072609 | Healthy Control |
| SRA | SRR5576291 | Healthy Control |
| ENA | ERZ267062 | Healthy Control |
| ENA | ERZ267021 | Healthy Control |
| ENA | ERZ267057 | Healthy Control |
| ENA | ERZ266973 | Healthy Control |
| ENA | ERZ487572 | Healthy Control |
| ENA | ERZ487610 | Healthy Control |
| ENA | ERZ376095 | Healthy Control |
| ENA | ERZ376154 | Healthy Control |
| ENA | ERZ376098 | Healthy Control |
| ENA | ERZ266997 | Healthy Control |
| ENA | ERZ267022 | Healthy Control |
| ENA | ERZ266976 | Healthy Control |
| ENA | ERZ267013 | Healthy Control |
| ENA | ERZ266983 | Healthy Control |
| ENA | ERZ487639 | Healthy Control |
| ENA | ERZ267048 | Healthy Control |
| ENA | ERZ267036 | Healthy Control |
| ENA | ERZ267025 | Healthy Control |
| ENA | ERZ487558 | Healthy Control |
| ENA | ERZ376132 | Healthy Control |
| ENA | ERZ376166 | Healthy Control |

| | | |
|---|---|---|
| ENA | ERZ266985 | Healthy Control |
| ENA | ERZ267063 | Healthy Control |
| ENA | ERZ376119 | Healthy Control |
| ENA | ERZ266998 | Healthy Control |
| ENA | ERZ376128 | Healthy Control |
| ENA | ERZ487601 | Healthy Control |
| ENA | ERZ487590 | Healthy Control |
| ENA | ERZ487594 | Healthy Control |
| ENA | ERZ376037 | Healthy Control |
| ENA | ERZ376162 | Healthy Control |
| ENA | ERZ266994 | Healthy Control |
| ENA | ERZ376115 | Healthy Control |
| ENA | ERZ267024 | Healthy Control |
| ENA | ERZ376051 | Healthy Control |
| ENA | ERZ376146 | Healthy Control |
| ENA | ERZ376039 | Healthy Control |
| ENA | ERZ266980 | Healthy Control |
| ENA | ERZ267061 | Healthy Control |
| ENA | ERZ376036 | Healthy Control |
| ENA | ERZ376116 | Healthy Control |
| ENA | ERZ267009 | Healthy Control |
| ENA | ERZ487644 | Healthy Control |
| ENA | ERZ267064 | Healthy Control |
| ENA | ERZ267018 | Healthy Control |
| ENA | ERZ376148 | Healthy Control |
| ENA | ERZ376147 | Healthy Control |
| ENA | ERZ487568 | Healthy Control |
| ENA | ERZ376032 | Healthy Control |
| ENA | ERZ376048 | Healthy Control |
| ENA | ERZ376045 | Healthy Control |
| ENA | ERZ487611 | Healthy Control |
| ENA | ERZ487642 | Healthy Control |
| ENA | ERZ487625 | Healthy Control |
| ENA | ERZ376133 | Healthy Control |
| ENA | ERZ376117 | Healthy Control |
| ENA | ERZ376058 | Healthy Control |
| ENA | ERZ487600 | Healthy Control |
| ENA | ERZ376040 | Healthy Control |
| ENA | ERZ376089 | Healthy Control |
| ENA | ERZ267002 | Healthy Control |
| ENA | ERZ376082 | Healthy Control |

| | | |
|---|---|---|
| ENA | ERZ267032 | Healthy Control |
| ENA | ERZ266995 | Healthy Control |
| ENA | ERZ266989 | Healthy Control |
| ENA | ERZ487634 | Healthy Control |
| ENA | ERZ376139 | Healthy Control |
| ENA | ERZ376102 | Healthy Control |
| ENA | ERZ487578 | Healthy Control |
| ENA | ERZ267054 | Healthy Control |
| ENA | ERZ266999 | Healthy Control |
| ENA | ERZ376136 | Healthy Control |
| ENA | ERZ376138 | Healthy Control |
| ENA | ERZ376092 | Healthy Control |
| ENA | ERZ376104 | Healthy Control |
| ENA | ERZ266974 | Healthy Control |
| ENA | ERZ267023 | Healthy Control |
| ENA | ERZ267033 | Healthy Control |
| ENA | ERZ267053 | Healthy Control |
| ENA | ERZ267055 | Healthy Control |
| ENA | ERZ376073 | Healthy Control |
| ENA | ERZ376055 | Healthy Control |
| ENA | ERZ376110 | Healthy Control |
| ENA | ERZ267066 | Healthy Control |
| ENA | ERZ267004 | Healthy Control |
| ENA | ERZ376152 | Healthy Control |
| ENA | ERZ487606 | Healthy Control |
| ENA | ERZ376050 | Healthy Control |

**Supplementary Table 3**. RNA-seq datasets used in this study. *RNA-seq datasets generated by this study

| Gene | Ensembl | Genome Position | logFC | Avg Expression | T Statistic | *P*-alue | FDR |
|---|---|---|---|---|---|---|---|
| GPRIN3 | ENSG00000185477 | chr4:90157537-90229161 | 0.155 | 4.478 | 4.918 | 1,14E-06 | 1,04E-03 |
| FAM65B | ENSG00000111913 | chr6:24797601-25042238 | 0.116 | 5.904 | 4.616 | 4,82E-06 | 2,73E-03 |
| CALB1 | ENSG00000104327 | chr8:91070836-91107703 | 0.218 | 5.241 | 4.528 | 7,24E-06 | 3,39E-03 |
| IGF2 | ENSG00000167244 | chr11:2150342-2170833 | -0.405 | 2.372 | -4.411 | 1,23E-05 | 4,68E-03 |
| AFF2 | ENSG00000155966 | chrX:147582139-148082193 | 0.116 | 5.72 | 4.291 | 2,09E-05 | 6,38E-03 |
| ARHGEF4 | ENSG00000136002 | chr2:131594489-131804836 | -0.114 | 7.001 | -4.153 | 3,77E-05 | 8,73E-03 |
| PHYHD1 | ENSG00000175287 | chr9:131683174-131704320 | -0.205 | 2.598 | -4.065 | 5,48E-05 | 1,02E-02 |
| SHD | ENSG00000105251 | chr19:4278598-4290721 | -0.208 | 2.058 | -3.974 | 7,98E-05 | 1,17E-02 |
| SAPCD2 | ENSG00000186193 | chr9:139956581-139965040 | -0.194 | 2.54 | -3.952 | 8,72E-05 | 1,26E-02 |
| TMEM196 | ENSG00000173452 | chr7:19758933-19813221 | 0.13 | 4.843 | 3.849 | 1,32E-04 | 1,57E-02 |
| HSPA12B | ENSG00000132622 | chr20:3713314-3733758 | -0.188 | 1.53 | -3.792 | 1,65E-04 | 1,68E-02 |
| SLC35F2 | ENSG00000110660 | chr11:107661717-107799019 | 0.118 | 2.891 | 3.729 | 2,11E-04 | 1,87E-02 |
| VAT1L | ENSG00000171724 | chr16:77822427-78014004 | 0.154 | 5.159 | 3.724 | 2,16E-04 | 1,89E-02 |
| . | ENSG00000255248 | chr11:121899063-121987031 | 0.098 | 6.977 | 3.542 | 4,30E-04 | 2,65E-02 |
| BCL6 | ENSG00000113916 | chr3:187439165-187463515 | 0.155 | 4.838 | 3.526 | 4,55E-04 | 2,73E-02 |
| CSRP1 | ENSG00000159176 | chr1:201452658-201478584 | -0.122 | 8.324 | -3.512 | 4,80E-04 | 2,82E-02 |
| CPLX1 | ENSG00000168993 | chr4:778745-819986 | -0.179 | 5.455 | -3.497 | 5,07E-04 | 2,87E-02 |
| ZC3HAV1 | ENSG00000105939 | chr7:138728266-138794465 | 0.152 | 4.309 | 3.484 | 5,32E-04 | 2,93E-02 |
| SLFN5 | ENSG00000166750 | chr17:33570055-33600674 | 0.113 | 4.824 | 3.476 | 5,47E-04 | 2,96E-02 |
| ZNF596 | ENSG00000172748 | chr8:182137-197342 | -0.124 | 3.252 | -3.478 | 5,44E-04 | 2,96E-02 |
| AK4 | ENSG00000162433 | chr1:65613232-65697828 | 0.099 | 6.338 | 3.448 | 6,06E-04 | 3,10E-02 |
| KNDC1 | ENSG00000171798 | chr10:134973951-135039916 | -0.131 | 6.392 | -3.443 | 6,18E-04 | 3,12E-02 |
| ADARB2 | ENSG00000185736 | chr10:1228073-1779670 | -0.152 | 3.735 | -3.417 | 6,77E-04 | 3,29E-02 |
| KAL1 | ENSG00000011201 | chrX:8496915-8700227 | 0.099 | 5.777 | 3.379 | 7,78E-04 | 3,48E-02 |
| NLRX1 | ENSG00000160703 | chr11:119037277-119054725 | -0.117 | 2.564 | -3.367 | 8,10E-04 | 3,53E-02 |
| NOL4 | ENSG00000101746 | chr18:31431064-31804916 | 0.071 | 5.95 | 3.32 | 9,58E-04 | 3,82E-02 |
| CNN1 | ENSG00000130176 | chr19:11649532-11661138 | -0.176 | 1.288 | -3.271 | 1,14E-03 | 4,19E-02 |
| HS3ST1 | ENSG00000002587 | chr4:11394774-11431389 | -0.099 | 2.917 | -3.258 | 1,19E-03 | 4,28E-02 |
| PTER | ENSG00000165983 | chr10:16478942-16555736 | 0.133 | 6.863 | 3.246 | 1,24E-03 | 4,40E-02 |
| SLC27A4 | ENSG00000167114 | chr9:131102925-131123749 | -0.09 | 4.877 | -3.229 | 1,31E-03 | 4,51E-02 |
| . | ENSG00000255571 | chr15:89904810-89939471 | -0.094 | 3.692 | -3.18 | 1,55E-03 | 4,81E-02 |
| COL5A1 | ENSG00000130635 | chr9:137533620-137736686 | -0.175 | 2.084 | -3.18 | 1,55E-03 | 4,81E-02 |
| ARL4D | ENSG00000175906 | chr17:41476327-41478505 | -0.167 | 2.834 | -3.148 | 1,73E-03 | 4,98E-02 |

**Supplementary Table 5.** Common DEGs in SZ NSC and DLPC. Data was obtained from SZDB 2.0. FDR<0.05 was considered as significant.

| Ch | start | end | width | strand | Fold | p.value | FDR | annotation | gwas snp id | gwas snp pos | gwas pvalue | gwas odds ratio | gwas region start | gwas region end | gwas_all_genes | distance to gwas snp | atac direction |
|---|---|---|---|---|---|---|---|---|---|---|---|---|---|---|---|---|---|
| chr17 | 5132397 | 5132842 | 446 | * | -2,07 | 1,45E-16 | 2,39E-11 | Exon (NM_001304284/USP6, exon 12 of 38) | rs2189336 | 5326162 | 4,52E-08 | 1,04477 | 4972162 | 5465662 | ZFP3,ZNF232,LOC101928000,USP6,ZNF594,LOC100130950,SCIMP,RABEP1,NUP88,RPAIN,C1QBP,DHX33,LOC105371506,DERL2,MIS12,LOC728392,NLRP1 | 193320 | Closed_in_SZ |
| chr17 | 78787512 | 78787997 | 486 | * | 1,71 | 4,02E-10 | 0,00000665 | Distal Intergenic | rs7225476 | 78561603 | 4,93E-08 | 0,96117 | 78406603 | 78788603 | ENDOV,NPTX1,RPTOR,LOC101928855 | 225909 | Open_in_SZ |
| chr6 | 26239964 | 26242003 | 2040 | * | 0,7 | 0,00000809 | 0,0216 | Promoter (<=1kb) | rs140365013 | 27523869 | 6,81E-41 | 1,22789 | 24933869 | 30563869 | RIPOR2,CMAHP,LOC101928663,CARMIL1,SCGN,HIST1H2AA,HIST1H2BA,HIST1H2APS1,SLC17A4,SLC17A1,SLC17A3,SLC17A2,TRIM38,HIST1H1A,HIST1H3A,HIST1H4A,HIST1H4B,HIST1H3B,HIST1H2AB,HIST1H2BB,HIST1H3C,HIST1H1C,LOC108783645,HFE,HIST1H4C,HIST1H1T,HIST1H2BC,HIST1H2AC,HIST1H1E,HIST1H2BD,HIST1H2BE,HIST1H4D,HIST1H3D,HIST1H2AD,HIST1H2BF,HIST1H4E,HIST1H2BG,HIST1H2AE,HIST1H3E,HIST1H1D,HIST1H4F,HIST1H4G,HIST1H3F,HIST1H2BH,HIST1H3G,HIST1H2BI,HIST1H4H,BTN3A2,BTN2A2,BTN3A1,BTN2A3P,BTN3A3,BTN2A1,LOC285819,BTN1A1,HCG11,HMGN4,LOC105374988,ABT1,ZNF322,GUSBP2,LINC00240,LOC100270746,HIST1H2BJ,HIST1H2AG,HIST1H2BK,HIST1H4I,HIST1H2AH,MIR3143,PRSS16,POM121L2,VN1R10P,ZNF204P,ZNF391,ZNF184,LINC01012,LOC100131289,HIST1H2BL,HIST1H2AI,HIST1H3H,HIST1H2AJ,HIST1H2BM,HIST1H4J,HIST1H4K,HIST1H2AK,HIST1H2BN,HIST1H2AL,HIST1H1B,HIST1H3I,HIST1H4L,HIST1H3J,HIST1H2AM,HIST1H2BO,OR2B2,OR2B6,ZNF165,ZSCAN12P1,ZSCAN16-AS1,ZSCAN16,ZKSCAN8,ZNF192P1,TOB2P1,ZSCAN9,ZKSCAN4,NKAPL,ZSCAN26,PGBD1,ZSCAN31,ZKSCAN3,ZSCAN12,ZSCAN23,GPX6,GPX5,ZBED9,LINC00533,LINC01623,HCG14,TRIM27,LINC01556,HCG15,ZNF311,LOC100129636,OR2W1,OR2B3,OR2J1,OR2J3,OR2J2,LOC101929006,OR14J1,OR5V1,OR12D3,OR12D2,OR12D1,OR11A1,OR10C1,OR2H1,MAS1L,LINC01015,UBD,SNORD32B,OR2H2,GABBR1,MOG,ZFP57,HLA-F,HLA-F-AS1,IFITM4P,HCG4,LOC554223,HLA-G,ZNRD1ASP,HCP5B,HLA-H,HCG4B,HLA-A,HCG9,HLA-J,ZNRD1,PPP1R11,RNF39,TRIM31,TRIM31-AS1,TRIM40,TRIM10,TRIM15,TRIM26,HCG17,HLA-L,MIR6891,HCG18,TRIM39,TRIM39-RPP21,RPP21,HLA-E,GNL1,PRR3,ABCF1,MIR877 | 1281866 | Open_in_SZ |
| chr11 | 104781102 | 104782982 | 1881 | * | 0,7 | 0,0000244 | 0,0465 | Distal Intergenic | rs1144403 | 104677870 | 3,36E-08 | 1,04144 | 104267870 | 104915870 | LINC02552,CASP12,LOC643733,CASP4,CASP5,CASP1,CARD16 | 103232 | Open_in_SZ |

**Supplementary Table 6.** Intersection of DARs with GWAS of schizophrenia patient samples.

| motif_id | motif_alt_id | sequence_name | start | stop | strand | score | p-value | q-value | matched_sequence |
|---|---|---|---|---|---|---|---|---|---|
| SP1_HUMAN.H11MO.0.A | | range=chr19 | 9498062 | 9498083 | - | 27,697 | 3,67E-10 | 0,00000046 | CGGGGGCGGGGCTGGAGGTGGG |
| SP3_HUMAN.H11MO.0.B | | range=chr19 | 9498067 | 9498086 | - | 21,8349 | 4,55E-09 | 0,00000566 | GCCCGGGGGCGGGGCTGGAG |
| KLF15_HUMAN.H11MO.0.A | | range=chr19 | 9498063 | 9498081 | - | 22,9091 | 7,32E-09 | 0,0000115 | GGGGCGGGGCTGGAGGTGG |
| SP2_HUMAN.H11MO.0.A | | range=chr19 | 9498066 | 9498087 | - | 22,5772 | 7,45E-09 | 0,00000811 | TGCCCGGGGGCGGGGCTGGAGG |
| RXRA_HUMAN.H11MO.0.A | | range=chr19 | 9498417 | 9498436 | + | 18,5806 | 2,11E-08 | 0,0000326 | GGGGGTGGCAGAGGTCACAA |
| MA0516.1 | SP2 | range=chr19 | 9498072 | 9498086 | + | 20,1207 | 2,54E-08 | 0,0000417 | GCCCCGCCCCCGGGC |
| KLF6_HUMAN.H11MO.0.A | | range=chr19 | 9498066 | 9498084 | - | 19,5606 | 3,88E-08 | 0,0000654 | CCGGGGGCGGGGCTGGAGG |
| KLF3_HUMAN.H11MO.0.B | | range=chr19 | 9498068 | 9498086 | - | 20,1515 | 4,07E-08 | 0,0000654 | GCCCGGGGGCGGGGCTGGA |
| ZNF76_HUMAN.H11MO.0.C | | range=chr19 | 9498563 | 9498584 | + | 20,374 | 7,13E-08 | 0,000108 | GGAGCCGGGTGGGAAGTGCAGT |
| MA0079.3 | SP1 | range=chr19 | 9498072 | 9498082 | + | 19,1538 | 7,65E-08 | 0,000128 | GCCCCGCCCCC |
| KLF9_HUMAN.H11MO.0.C | | range=chr19 | 9498250 | 9498264 | - | 20,4286 | 8,09E-08 | 0,000115 | GGAAGGGGCGTGGCT |
| MAZ_HUMAN.H11MO.0.A | | range=chr19 | 9498061 | 9498082 | - | 19,662 | 9,04E-08 | 0,000132 | GGGGGCGGGGCTGGAGGTGGGA |
| PATZ1_HUMAN.H11MO.0.C | | range=chr19 | 9498061 | 9498082 | - | 19,3472 | 9,11E-08 | 0,000127 | GGGGGCGGGGCTGGAGGTGGGA |
| MA0810.1 | TFAP2A(var.2) | range=chr19 | 9498076 | 9498087 | + | 16,8364 | 1,34E-07 | 0,000169 | CGCCCCCGGGCA |
| SP2_HUMAN.H11MO.0.A | | range=chr19 | 9498085 | 9498106 | - | 19,3252 | 1,36E-07 | 0,0000742 | GGGCTGTGGGCGGGGCCCATGC |
| MA0499.1 | Myod1 | range=chr19 | 9498815 | 9498827 | - | 16,5918 | 1,42E-07 | 0,000221 | AGCAGCTGCCCCC |
| MA0741.1 | KLF16 | range=chr19 | 9498072 | 9498082 | + | 18,2414 | 0,00000017 | 0,00026 | GCCCCGCCCCC |
| PATZ1_HUMAN.H11MO.0.C | | range=chr19 | 9498066 | 9498087 | - | 18,3958 | 2,12E-07 | 0,000148 | TGCCCGGGGGCGGGGCTGGAGG |
| HEN1_HUMAN.H11MO.0.C | | range=chr19 | 9498813 | 9498832 | + | 18,9634 | 2,15E-07 | 0,000327 | AAGGGGGCAGCTGCTTCCTT |
| SP1_HUMAN.H11MO.0.A | | range=chr19 | 9498081 | 9498102 | - | 18,1061 | 2,28E-07 | 0,000143 | TGTGGGCGGGGCCCATGCCCGG |
| KLF12_HUMAN.H11MO.0.C | | range=chr19 | 9498072 | 9498082 | - | 19,2121 | 2,46E-07 | 0,000402 | GGGGGCGGGGC |
| KLF9_HUMAN.H11MO.0.C | | range=chr19 | 9498090 | 9498104 | - | 19,0857 | 2,51E-07 | 0,000178 | GCTGTGGGCGGGGCC |
| SP4_HUMAN.H11MO.0.A | | range=chr19 | 9498068 | 9498087 | - | 18,1048 | 2,53E-07 | 0,000397 | TGCCCGGGGGCGGGGCTGGA |
| WT1_HUMAN.H11MO.0.C | | range=chr19 | 9498067 | 9498086 | - | 18,6364 | 2,71E-07 | 0,000351 | GCCCGGGGGCGGGGCTGGAG |
| KLF1_HUMAN.H11MO.0.A | | range=chr19 | 9498068 | 9498081 | - | 18,3636 | 2,75E-07 | 0,000456 | GGGGCGGGGCTGGA |
| MA0599.1 | KLF5 | range=chr19 | 9498072 | 9498081 | + | 17 | 3,39E-07 | 0,000571 | GCCCCGCCCC |
| AP2B_HUMAN.H11MO.0.B | | range=chr19 | 9498077 | 9498086 | - | 16,0112 | 3,39E-07 | 0,000511 | GCCCGGGGGC |
| MA0740.1 | KLF14 | range=chr19 | 9498250 | 9498263 | + | 18,6724 | 3,59E-07 | 0,000558 | AGCCACGCCCCTTC |
| MA0685.1 | SP4 | range=chr19 | 9498248 | 9498264 | + | 18,4 | 3,79E-07 | 0,000602 | CGAGCCACGCCCCTTCC |
| MA0524.2 | TFAP2C | range=chr19 | 9498076 | 9498087 | + | 16,2959 | 4,33E-07 | 0,000598 | CGCCCCCGGGCA |
| MA0811.1 | TFAP2B | range=chr19 | 9498076 | 9498087 | + | 16,3793 | 4,38E-07 | 0,000618 | CGCCCCCGGGCA |
| PO5F1_HUMAN.H11MO.0.A | | range=chr19 | 9498829 | 9498844 | + | 17,3818 | 4,53E-07 | 0,000819 | CCTTTTTTATGCAAAC |
| OLIG2_HUMAN.H11MO.0.B | | range=chr19 | 9498819 | 9498836 | + | 16,3333 | 4,82E-07 | 0,000787 | GCAGCTGCTTCCTTTTTT |
| INSM1_HUMAN.H11MO.0.C | | range=chr19 | 9498810 | 9498821 | + | 18,6327 | 5,33E-07 | 0,000876 | TTTAAGGGGGCA |
| SP4_HUMAN.H11MO.0.A | | range=chr19 | 9498056 | 9498075 | - | 17,0645 | 6,42E-07 | 0,000503 | GGGCTGGAGGTGGGACCAGA |

| | | | | | | | | | |
|---|---|---|---|---|---|---|---|---|---|
| MA0746.1 | SP3 | range=chr19 | 9498072 | 9498082 | + | 15,9837 | 7,41E-07 | 0,000667 | GCCCCGCCCCC |
| MA0516.1 | SP2 | range=chr19 | 9498091 | 9498105 | + | 17,1724 | 7,83E-07 | 0,000641 | GCCCCGCCCACAGCC |
| SP3_HUMAN.H11MO.0.B | | range=chr19 | 9498086 | 9498105 | - | 16,7248 | 9,02E-07 | 0,00056 | GGCTGTGGGCGGGGCCCATG |
| HEN1_HUMAN.H11MO.0.C | | range=chr19 | 9498813 | 9498832 | - | 16,5244 | 9,05E-07 | 0,000689 | AAGGAAGCAGCTGCCCCCTT |
| KLF12_HUMAN.H11MO.0.C | | range=chr19 | 9498251 | 9498261 | - | 17,5758 | 9,39E-07 | 0,000767 | AGGGGCGTGGC |
| KLF9_HUMAN.H11MO.0.C | | range=chr19 | 9498071 | 9498085 | - | 17,2 | 9,49E-07 | 0,000448 | CCCGGGGGCGGGGCT |
| SP2_HUMAN.H11MO.0.A | | range=chr19 | 9498061 | 9498082 | - | 16,6911 | 0,000001 | 0,000321 | GGGGGCGGGGCTGGAGGTGGGA |
| MA0741.1 | KLF16 | range=chr19 | 9498251 | 9498261 | + | 16,8621 | 0,00000104 | 0,000793 | GCCACGCCCCT |
| MA0746.1 | SP3 | range=chr19 | 9498251 | 9498261 | + | 15,6748 | 0,00000104 | 0,000667 | GCCACGCCCCT |
| SP2_HUMAN.H11MO.0.A | | range=chr19 | 9498054 | 9498075 | - | 16,4553 | 0,00000118 | 0,000321 | GGGCTGGAGGTGGGACCAGACC |
| MA0746.1 | SP3 | range=chr19 | 9498091 | 9498101 | + | 15,6341 | 0,00000125 | 0,000667 | GCCCCGCCCAC |
| SP4_HUMAN.H11MO.0.A | | range=chr19 | 9498087 | 9498106 | - | 16,2016 | 0,00000132 | 0,000689 | GGGCTGTGGGCGGGGCCCAT |
| PATZ1_HUMAN.H11MO.0.C | | range=chr19 | 9498060 | 9498081 | - | 16,0833 | 0,00000139 | 0,000649 | GGGGCGGGGCTGGAGGTGGGAC |
| KLF12_HUMAN.H11MO.0.C | | range=chr19 | 9498091 | 9498101 | - | 16,8636 | 0,00000144 | 0,000787 | GTGGGCGGGGC |
| MA0741.1 | KLF16 | range=chr19 | 9498091 | 9498101 | + | 16,5345 | 0,00000155 | 0,000793 | GCCCCGCCCAC |

**Supplementary Table 7.** FIMO analysis of the regulatory region at ZNF560 promoter

| Rank | Motif (logo) | P-value | log P-pvalue | % of Targets | % of Background | STD(Bg STD) | Best Match/Details |
|---|---|---|---|---|---|---|---|
| 1 | 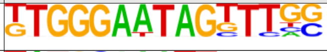 | 1e-15 | -3.592e+01 | 0.28% | 0.00% | 67.8bp (21.2bp) | RBPJ/MA1116.1/Jaspar(0.670) More Information \| Similar Motifs Found |
| 2 | 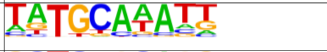 | 1e-14 | -3.290e+01 | 14.61% | 9.69% | 56.6bp (64.3bp) | POU2F3/MA0627.2/Jaspar(0.930) More Information \| Similar Motifs Found |
| 3 | 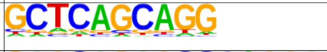 | 1e-14 | -3.250e+01 | 27.64% | 21.10% | 53.4bp (60.2bp) | Zic(Zf)/Cerebellum-ZIC1.2-ChIP-Seq(GSE60731)/Homer(0.886) More Information \| Similar Motifs Found |
| 4 | 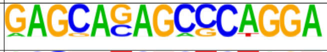 | 1e-13 | -3.199e+01 | 0.36% | 0.01% | 35.6bp (43.6bp) | HNF4a(NR),DR1/HepG2-HNF4a-ChIP-Seq(GSE25021)/Homer(0.561) More Information \| Similar Motifs Found |
| 5 | 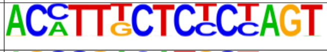 | 1e-13 | -3.006e+01 | 0.24% | 0.00% | 49.8bp (0.0bp) | PRDM1/MA0508.3/Jaspar(0.622) More Information \| Similar Motifs Found |
| 6 | 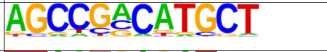 | 1e-12 | -2.799e+01 | 0.97% | 0.14% | 53.0bp (57.6bp) | YY1(Zf)/Promoter/Homer(0.658) More Information \| Similar Motifs Found |
| 7 | 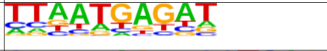 | 1e-11 | -2.636e+01 | 12.14% | 8.11% | 55.1bp (63.7bp) | PH0151.1_Pou6f1_1/Jaspar(0.834) More Information \| Similar Motifs Found |
| 8 | 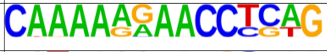 | 1e-11 | -2.627e+01 | 0.28% | 0.00% | 52.8bp (24.5bp) | ZNF384/MA1125.1/Jaspar(0.639) More Information \| Similar Motifs Found |
| 9 | 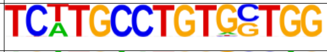 | 1e-11 | -2.592e+01 | 0.24% | 0.00% | 28.7bp (9.7bp) | SD0002.1_at_AC_acceptor/Jaspar(0.621) More Information \| Similar Motifs Found |
| 10 | 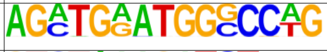 | 1e-11 | -2.592e+01 | 0.24% | 0.00% | 40.8bp (22.6bp) | Hoxb4(Homeobox)/ES-Hoxb4-ChIP-Seq(GSE34014)/Homer(0.606) More Information \| Similar Motifs Found |
| 11 | 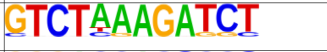 | 1e-11 | -2.586e+01 | 0.36% | 0.01% | 54.4bp (50.9bp) | HNF4A(var.2)/MA1494.1/Jaspar(0.633) More Information \| Similar Motifs Found |
| 12 | 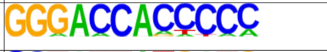 | 1e-11 | -2.580e+01 | 0.65% | 0.06% | 44.1bp (50.8bp) | GLI3(Zf)/Limb-GLI3-ChIP-Chip(GSE11077)/Homer(0.791) More Information \| Similar Motifs Found |
| 13 | 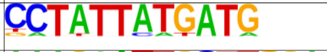 | 1e-11 | -2.540e+01 | 0.32% | 0.01% | 50.1bp (54.1bp) | PB0174.1_Sox30_2/Jaspar(0.708) More Information \| Similar Motifs Found |
| 14 | 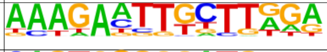 | 1e-10 | -2.477e+01 | 1.05% | 0.20% | 54.7bp (61.3bp) | PB0175.1_Sox4_2/Jaspar(0.609) More Information \| Similar Motifs Found |
| 15 | 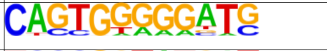 | 1e-10 | -2.438e+01 | 0.89% | 0.14% | 61.7bp (60.0bp) | Znf281/MA1630.1/Jaspar(0.727) More Information \| Similar Motifs Found |
| 16 | 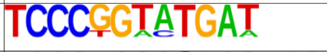 | 1e-10 | -2.436e+01 | 0.20% | 0.00% | 60.7bp (0.0bp) | PB0140.1_Irf6_2/Jaspar(0.617) More Information \| Similar Motifs Found |
| 17 | 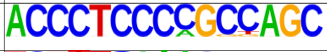 | 1e-10 | -2.436e+01 | 0.20% | 0.00% | 66.6bp (0.0bp) | PB0164.1_Smad3_2/Jaspar(0.630) More Information \| Similar Motifs Found |
| 18 | 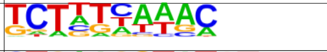 | 1e-9 | -2.268e+01 | 2.19% | 0.80% | 58.5bp (60.2bp) | CHR(?)/Hela-CellCycle-Expression/Homer(0.615) More Information \| Similar Motifs Found |
| 19 | 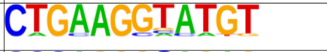 | 1e-9 | -2.206e+01 | 0.45% | 0.03% | 65.0bp (67.3bp) | SD0001.1_at_AC_acceptor/Jaspar(0.655) More Information \| Similar Motifs Found |
| 20 | 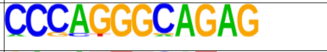 | 1e-9 | -2.198e+01 | 0.77% | 0.12% | 55.7bp (59.0bp) | EBF1/MA0154.4/Jaspar(0.687) More Information \| Similar Motifs Found |
| 21 | 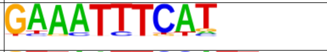 | 1e-9 | -2.196e+01 | 2.55% | 1.04% | 57.7bp (63.4bp) | PB0169.1_Sox15_2/Jaspar(0.739) More Information \| Similar Motifs Found |
| 22 | 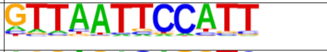 | 1e-9 | -2.178e+01 | 1.82% | 0.61% | 51.2bp (64.0bp) | Prop1(Homeobox)/GHFT1-PROP1.biotin-ChIP-Seq(GSE77302)/Homer(0.759) More Information \| Similar Motifs Found |
| 23 | 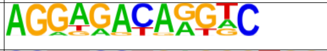 | 1e-9 | -2.134e+01 | 1.25% | 0.33% | 49.0bp (58.8bp) | RXR(NR),DR1/3T3L1-RXR-ChIP-Seq(GSE13511)/Homer(0.646) More Information \| Similar Motifs Found |
| 24 | 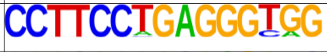 | 1e-9 | -2.091e+01 | 0.20% | 0.00% | 42.0bp (26.0bp) | ETV4/MA0764.2/Jaspar(0.643) More Information \| Similar Motifs Found |
| 25 | 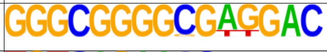 | 1e-9 | -2.091e+01 | 0.20% | 0.00% | 46.5bp (24.6bp) | KLF15/MA1513.1/Jaspar(0.752) More Information \| Similar Motifs Found |
| 26 | 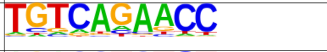 | 1e-9 | -2.091e+01 | 5.71% | 3.31% | 55.3bp (60.4bp) | Tbx20(T-box)/Heart-Tbx20-ChIP-Seq(GSE29636)/Homer(0.693) More Information \| Similar Motifs Found |
| 27 | 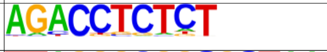 | 1e-9 | -2.078e+01 | 2.67% | 1.15% | 52.6bp (60.6bp) | ZNF768(Zf)/Rajj-ZNF768-ChIP-Seq(GSE111879)/Homer(0.683) More Information \| Similar Motifs Found |
| 28 | 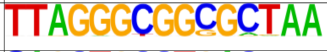 | 1e-8 | -2.047e+01 | 0.24% | 0.00% | 40.7bp (36.2bp) | PB0039.1_Klf7_1/Jaspar(0.637) More Information \| Similar Motifs Found |
| 29 | 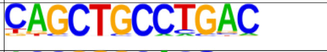 | 1e-8 | -2.035e+01 | 0.85% | 0.16% | 51.0bp (59.3bp) | ASCL1/MA1100.2/Jaspar(0.722) More Information \| Similar Motifs Found |
| 30 | 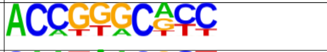 | 1e-8 | -1.973e+01 | 1.09% | 0.27% | 46.7bp (55.3bp) | ZBTB14/MA1650.1/Jaspar(0.611) More Information \| Similar Motifs Found |
| 31 | 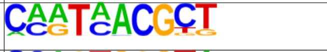 | 1e-8 | -1.930e+01 | 0.40% | 0.03% | 61.6bp (50.9bp) | OVOL1/MA1544.1/Jaspar(0.682) More Information \| Similar Motifs Found |
| 32 | 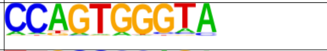 | 1e-8 | -1.925e+01 | 2.27% | 0.94% | 61.2bp (61.7bp) | ZNF75D/MA1601.1/Jaspar(0.646) More Information \| Similar Motifs Found |
| 33 | 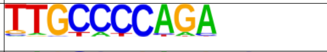 | 1e-8 | -1.905e+01 | 2.71% | 1.23% | 51.2bp (61.2bp) | Plagl1/MA1615.1/Jaspar(0.746) More Information \| Similar Motifs Found |
| 34 | 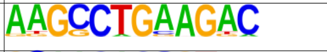 | 1e-8 | -1.867e+01 | 0.53% | 0.06% | 59.9bp (57.5bp) | ELF1(ETS)/Jurkat-ELF1-ChIP-Seq(SRA014231)/Homer(0.595) More Information \| Similar Motifs Found |
| 35 | 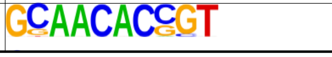 | 1e-8 | -1.858e+01 | 0.73% | 0.13% | 56.2bp (59.5bp) | PB0117.1_Eomes_2/Jaspar(0.642) More Information \| Similar Motifs Found |

| | | | | | | | |
|---|---|---|---|---|---|---|---|
| 36 | CGTGTGGGCG | 1e-7 | -1.819e+01 | 1.66% | 0.60% | 57.1bp (54.2bp) | Egr2(Zf)/Thymocytes-Egr2-ChIP-Seq(GSE34254)/Homer(0.872) More Information \| Similar Motifs Found |
| 37 | CCTAAGCATCGC | 1e-7 | -1.748e+01 | 0.20% | 0.00% | 61.0bp (45.7bp) | ZNF460/MA1596.1/Jaspar(0.577) More Information \| Similar Motifs Found |
| 38 | TGCGAGGGGGCAGGA | 1e-7 | -1.701e+01 | 0.24% | 0.01% | 57.5bp (64.1bp) | INSM1/MA0155.1/Jaspar(0.753) More Information \| Similar Motifs Found |
| 39 | TGGCGTGG | 1e-7 | -1.673e+01 | 10.08% | 7.15% | 56.2bp (56.5bp) | Ahr::Arnt/MA0006.1/Jaspar(0.849) More Information \| Similar Motifs Found |
| 40 | TTGTTGTTCT | 1e-7 | -1.648e+01 | 1.70% | 0.66% | 53.6bp (62.2bp) | PB0122.1_Foxk1_2/Jaspar(0.750) More Information \| Similar Motifs Found |
| 41 | CCCACAGAGAATGGC | 1e-7 | -1.617e+01 | 0.36% | 0.03% | 55.0bp (58.9bp) | RUNX1/MA0002.2/Jaspar(0.594) More Information \| Similar Motifs Found |
| 42 | TGGTGCCCTCTGTAA | 1e-6 | -1.608e+01 | 0.16% | 0.00% | 37.6bp (1.7bp) | Erra(NR)/HepG2-Erra-ChIP-Seq(GSE31477)/Homer(0.633) More Information \| Similar Motifs Found |
| 43 | CCCCGTGGTGCCAGG | 1e-6 | -1.608e+01 | 0.16% | 0.00% | 11.3bp (31.2bp) | Plagl1/MA1615.1/Jaspar(0.614) More Information \| Similar Motifs Found |
| 44 | CCCTCACCTCTGCAA | 1e-6 | -1.573e+01 | 0.32% | 0.02% | 58.0bp (55.6bp) | ZKSCAN5/MA1652.1/Jaspar(0.624) More Information \| Similar Motifs Found |
| 45 | TGGAGGCGAG | 1e-6 | -1.565e+01 | 3.72% | 2.08% | 56.2bp (58.5bp) | ZKSCAN5/MA1652.1/Jaspar(0.733) More Information \| Similar Motifs Found |
| 46 | AGGCTTGGATGG | 1e-6 | -1.514e+01 | 0.40% | 0.04% | 57.6bp (65.7bp) | Pax8(Paired,Homeobox)/Thyroid-Pax8-ChIP-Seq(GSE26938)/Homer(0.672) More Information \| Similar Motifs Found |
| 47 | TGTGAACTTA | 1e-6 | -1.493e+01 | 0.73% | 0.17% | 59.6bp (62.4bp) | VDR/MA0693.2/Jaspar(0.786) More Information \| Similar Motifs Found |
| 48 | CCACCTCCCGTC | 1e-6 | -1.445e+01 | 0.24% | 0.01% | 31.9bp (56.2bp) | ETS:RUNX(ETS,Runt)/Jurkat-RUNX1-ChIP-Seq(GSE17954)/Homer(0.650) More Information \| Similar Motifs Found |
| 49 | CCAGCCCGCCCATTC | 1e-6 | -1.408e+01 | 0.20% | 0.01% | 45.8bp (46.7bp) | PB0164.1_Smad3_2/Jaspar(0.624) More Information \| Similar Motifs Found |
| 50 | CCCTAACAGGGACCC | 1e-5 | -1.354e+01 | 0.12% | 0.00% | 61.1bp (0.0bp) | PLAGL2/MA1548.1/Jaspar(0.584) More Information \| Similar Motifs Found |
| 51 | AGGTGCTGTGGGTGG | 1e-5 | -1.354e+01 | 0.12% | 0.00% | 54.5bp (0.0bp) | TCF4/MA0830.2/Jaspar(0.602) More Information \| Similar Motifs Found |
| 52 | AGCTCTGATTCCGCC | 1e-5 | -1.354e+01 | 0.12% | 0.00% | 51.2bp (0.0bp) | Gfi1b(Zf)/HPC7-Gfi1b-ChIP-Seq(GSE22178)/Homer(0.634) More Information \| Similar Motifs Found |
| 53 | AACAAGCACC | 1e-5 | -1.297e+01 | 3.89% | 2.35% | 54.6bp (62.1bp) | MF0011.1_HMG_class/Jaspar(0.608) More Information \| Similar Motifs Found |
| 54 | ATAGTAATTA | 1e-5 | -1.205e+01 | 19.63% | 16.27% | 55.1bp (65.0bp) | Dlx3(Homeobox)/Kerainocytes-Dlx3-ChIP-Seq(GSE89884)/Homer(0.806) More Information \| Similar Motifs Found |
| 55 | TCATTCCCAGCCACG | 1e-4 | -1.073e+01 | 0.24% | 0.02% | 43.7bp (58.8bp) | Zfp809(Zf)/ES-Zfp809-ChIP-Seq(GSE70799)/Homer(0.652) More Information \| Similar Motifs Found |
| 56 | AGTTCACCTCAG | 1e-4 | -9.934e+00 | 0.20% | 0.02% | 57.1bp (51.5bp) | NR1I3/MA1534.1/Jaspar(0.671) More Information \| Similar Motifs Found |
| 57 | GTAATTCCAGAG | 1e-4 | -9.423e+00 | 0.12% | 0.00% | 46.9bp (40.5bp) | Dlx3(Homeobox)/Kerainocytes-Dlx3-ChIP-Seq(GSE89884)/Homer(0.662) More Information \| Similar Motifs Found |
| 58 | GGGGCAGAGAGAGGC | 1e-1 | -3.920e+00 | 0.04% | 0.00% | 35.9bp (0.0bp) | PB0202.1_Zfp410_2/Jaspar(0.601) More Information \| Similar Motifs Found |
| 59 | CCTCCCACCCTTGCC | 1e-1 | -3.920e+00 | 0.04% | 0.00% | 51.8bp (0.0bp) | Egr2(Zf)/Thymocytes-Egr2-ChIP-Seq(GSE34254)/Homer(0.649) More Information \| Similar Motifs Found |
| 60 | CCCGTGATCCAGCCA | 1e-1 | -3.920e+00 | 0.04% | 0.00% | 27.8bp (0.0bp) | TFE3/MA0831.2/Jaspar(0.581) More Information \| Similar Motifs Found |

**Supplementary Table 8.** Motif enrichment analysis of regions with increased accessibility in SZ iPSC.

| Rank | Motif (logo) | P-value | log P-pvalue | % of Targets | % of Background | STD(Bg STD) | Best Match/Details |
|---|---|---|---|---|---|---|---|
| 1 | | 1e-14 | -3.443e+01 | 0.53% | 0.00% | 55.7bp (28.1bp) | USF2/MA0526.3/Jaspar(0.728) More Information \| Similar Motifs Found |
| 2 | | 1e-14 | -3.274e+01 | 0.39% | 0.00% | 65.0bp (78.6bp) | PPARE(NR),DR1/3T3L1-Pparg-ChIP-Seq(GSE13511)/Homer(0.657) More Information \| Similar Motifs Found |
| 3 | | 1e-13 | -3.027e+01 | 1.84% | 0.30% | 54.2bp (54.8bp) | PB0195.1_Zbtb3_2/Jaspar(0.698) More Information \| Similar Motifs Found |
| 4 | | 1e-12 | -2.859e+01 | 0.39% | 0.00% | 59.5bp (10.6bp) | NFIC::TLX1/MA0119.1/Jaspar(0.579) More Information \| Similar Motifs Found |
| 5 | | 1e-12 | -2.771e+01 | 1.25% | 0.14% | 57.9bp (50.6bp) | ETS:RUNX(ETS,Runt)/Jurkat-RUNX1-ChIP-Seq(GSE17954)/Homer(0.613) More Information \| Similar Motifs Found |
| 6 | | 1e-12 | -2.765e+01 | 19.66% | 13.16% | 51.7bp (55.1bp) | CTCF/MA0139.1/Jaspar(0.718) More Information \| Similar Motifs Found |
| 7 | | 1e-11 | -2.726e+01 | 0.59% | 0.01% | 55.0bp (56.5bp) | E2F6/MA0471.2/Jaspar(0.598) More Information \| Similar Motifs Found |
| 8 | | 1e-11 | -2.617e+01 | 0.39% | 0.00% | 49.7bp (88.7bp) | Srebp2(bHLH)/HepG2-Srebp2-ChIP-Seq(GSE31477)/Homer(0.538) More Information \| Similar Motifs Found |
| 9 | | 1e-11 | -2.611e+01 | 0.85% | 0.05% | 57.7bp (56.1bp) | PB0025.1_Glis2_1/Jaspar(0.653) More Information \| Similar Motifs Found |
| 10 | | 1e-10 | -2.512e+01 | 1.45% | 0.22% | 54.8bp (55.1bp) | HIC1(Zf)/Treg-ZBTB29-ChIP-Seq(GSE99889)/Homer(0.647) More Information \| Similar Motifs Found |
| 11 | | 1e-10 | -2.508e+01 | 0.72% | 0.04% | 40.4bp (56.6bp) | NFYB/MA0502.2/Jaspar(0.680) More Information \| Similar Motifs Found |
| 12 | | 1e-10 | -2.398e+01 | 0.53% | 0.01% | 42.8bp (51.8bp) | GLI3(Zf)/Limb-GLI3-ChIP-Chip(GSE11077)/Homer(0.670) More Information \| Similar Motifs Found |
| 13 | | 1e-10 | -2.312e+01 | 0.39% | 0.00% | 49.4bp (42.4bp) | IRF1(IRF)/PBMC-IRF1-ChIP-Seq(GSE43036)/Homer(0.702) More Information \| Similar Motifs Found |
| 14 | | 1e-9 | -2.206e+01 | 0.59% | 0.03% | 52.1bp (58.6bp) | ZEB2(Zf)/SNU398-ZEB2-ChIP-Seq(GSE103048)/Homer(0.632) More Information \| Similar Motifs Found |
| 15 | | 1e-9 | -2.143e+01 | 1.97% | 0.50% | 55.7bp (52.4bp) | Nr2e1/MA0676.1/Jaspar(0.728) More Information \| Similar Motifs Found |
| 16 | | 1e-9 | -2.125e+01 | 4.54% | 1.98% | 55.1bp (57.9bp) | ZNF415(Zf)/HEK293-ZNF415.GFP-ChIP-Seq(GSE58341)/Homer(0.604) More Information \| Similar Motifs Found |
| 17 | | 1e-9 | -2.113e+01 | 0.39% | 0.01% | 24.6bp (41.6bp) | PB0130.1_Gm397_2/Jaspar(0.632) More Information \| Similar Motifs Found |
| 18 | | 1e-9 | -2.111e+01 | 0.33% | 0.00% | 47.4bp (56.6bp) | Foxq1/MA0040.1/Jaspar(0.676) More Information \| Similar Motifs Found |
| 19 | | 1e-9 | -2.099e+01 | 0.53% | 0.02% | 70.8bp (52.2bp) | PB0169.1_Sox15_2/Jaspar(0.703) More Information \| Similar Motifs Found |
| 20 | | 1e-9 | -2.090e+01 | 35.57% | 28.42% | 54.7bp (57.9bp) | NFIX/MA0671.1/Jaspar(0.619) More Information \| Similar Motifs Found |
| 21 | | 1e-9 | -2.084e+01 | 9.20% | 5.37% | 55.7bp (57.8bp) | FXR(NR),IR1/Liver-FXR-ChIP-Seq(Chong_et_al.)/Homer(0.642) More Information \| Similar Motifs Found |
| 22 | | 1e-8 | -2.062e+01 | 0.26% | 0.00% | 45.2bp (0.0bp) | TFAP2E/MA1569.1/Jaspar(0.564) More Information \| Similar Motifs Found |
| 23 | | 1e-8 | -2.062e+01 | 0.26% | 0.00% | 50.2bp (0.0bp) | ETV4/MA0764.2/Jaspar(0.657) More Information \| Similar Motifs Found |
| 24 | | 1e-8 | -2.054e+01 | 2.83% | 0.97% | 45.9bp (47.6bp) | ZNF384/MA1125.1/Jaspar(0.840) More Information \| Similar Motifs Found |
| 25 | | 1e-8 | -2.050e+01 | 1.25% | 0.21% | 50.5bp (50.5bp) | Arid3b/MA0601.1/Jaspar(0.697) More Information \| Similar Motifs Found |
| 26 | | 1e-8 | -2.043e+01 | 0.59% | 0.03% | 53.4bp (48.9bp) | PB0091.1_Zbtb3_1/Jaspar(0.536) More Information \| Similar Motifs Found |
| 27 | | 1e-8 | -2.002e+01 | 0.99% | 0.13% | 49.8bp (56.3bp) | Zfx/MA0146.2/Jaspar(0.644) More Information \| Similar Motifs Found |
| 28 | | 1e-8 | -1.969e+01 | 0.33% | 0.00% | 60.8bp (35.6bp) | ZNF415(Zf)/HEK293-ZNF415.GFP-ChIP-Seq(GSE58341)/Homer(0.679) More Information \| Similar Motifs Found |
| 29 | | 1e-8 | -1.964e+01 | 0.39% | 0.01% | 58.6bp (50.7bp) | Srebp1a(bHLH)/HepG2-Srebp1a-ChIP-Seq(GSE31477)/Homer(0.639) More Information \| Similar Motifs Found |
| 30 | | 1e-8 | -1.886e+01 | 0.53% | 0.02% | 46.8bp (50.8bp) | MGA/MA0801.1/Jaspar(0.717) More Information \| Similar Motifs Found |
| 31 | | 1e-7 | -1.823e+01 | 3.48% | 1.45% | 50.8bp (56.1bp) | TFAP2A(var.2)/MA0810.1/Jaspar(0.707) More Information \| Similar Motifs Found |
| 32 | | 1e-7 | -1.786e+01 | 0.92% | 0.13% | 42.9bp (57.5bp) | Zic1::Zic2/MA1628.1/Jaspar(0.685) More Information \| Similar Motifs Found |
| 33 | | 1e-7 | -1.785e+01 | 0.26% | 0.00% | 60.5bp (44.9bp) | PRDM1/MA0508.3/Jaspar(0.588) More Information \| Similar Motifs Found |
| 34 | | 1e-7 | -1.767e+01 | 2.30% | 0.77% | 48.6bp (60.6bp) | PB0164.1_Smad3_2/Jaspar(0.660) More Information \| Similar Motifs Found |
| 35 | | 1e-7 | -1.755e+01 | 4.34% | 2.05% | 54.6bp (55.8bp) | Dmrt1/MA1603.1/Jaspar(0.618) More Information \| Similar Motifs Found |
| 36 | | 1e-7 | -1.697e+01 | 2.50% | 0.90% | 56.7bp (57.7bp) | PB0094.1_Zfp128_1/Jaspar(0.730) More Information \| Similar Motifs Found |
| 37 | | 1e-7 | -1.647e+01 | 1.05% | 0.19% | 64.2bp (61.9bp) | SP3/MA0746.2/Jaspar(0.631) More Information \| Similar Motifs Found |
| 38 | | 1e-6 | -1.567e+01 | 0.72% | 0.09% | 44.7bp (55.4bp) | PB0008.1_E2F2_1/Jaspar(0.665) More Information \| Similar Motifs Found |
| 39 | | 1e-6 | -1.487e+01 | 0.20% | 0.00% | 57.1bp (0.0bp) | IRF8(IRF)/BMDM-IRF8-ChIP-Seq(GSE77884)/Homer(0.649) More Information \| Similar Motifs Found |
| 40 | | 1e-6 | -1.400e+01 | 0.79% | 0.13% | 58.8bp (54.1bp) | ZEB2(Zf)/SNU398-ZEB2-ChIP-Seq(GSE103048)/Homer(0.736) More Information \| Similar Motifs Found |
| 41 | | 1e-6 | -1.389e+01 | 0.33% | 0.01% | 28.8bp (48.2bp) | Sp1(Zf)/Promoter/Homer(0.650) More Information \| Similar Motifs Found |
| 42 | | 1e-5 | -1.280e+01 | 0.20% | 0.00% | 18.6bp (29.9bp) | RXRB(var.2)/MA1555.1/Jaspar(0.644) More Information \| Similar Motifs Found |
| 43 | | 1e-3 | -7.234e+00 | 0.13% | 0.00% | 52.9bp (21.0bp) | ZNF263/MA0528.2/Jaspar(0.716) More Information \| Similar Motifs Found |
| 44 | | 1e-1 | -4.362e+00 | 0.07% | 0.00% | 0.0bp (0.0bp) | KLF9/MA1107.2/Jaspar(0.833) More Information \| Similar Motifs Found |

**Supplementary Table 9.** Motif enrichment analysis of regions with increased accessibility in HC iPSC.

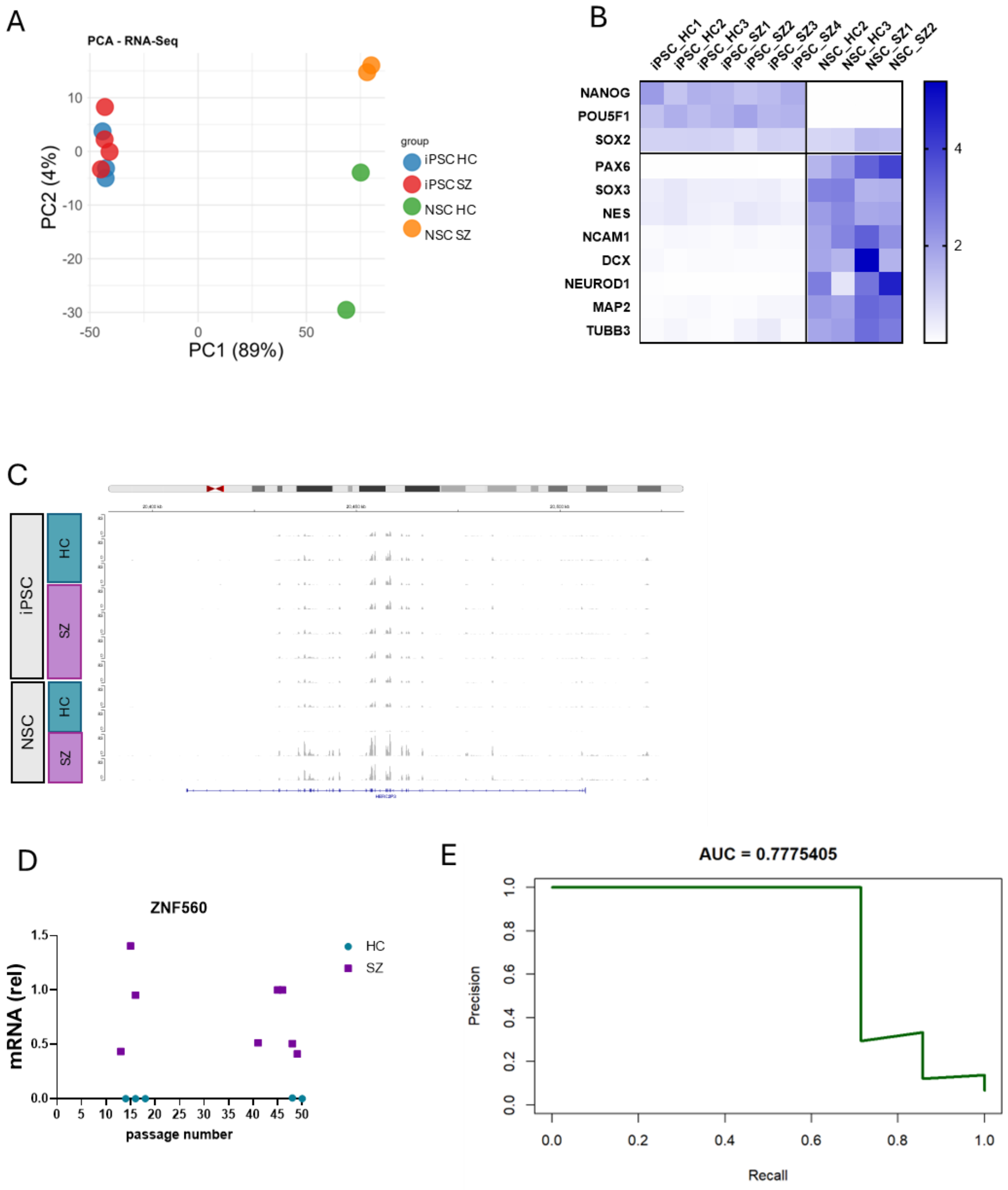


**Supplementary Figure 1**. **A.** Principal Component Analysis of RNA-seq data from iPSC and NSC. **B.** Heatmap of RNA-seq data from iPSC and NSC (both HC and SZ), showing classic pluripotency and neural genes. Data is mean-normalized (mean-centering). **C.** Visualization of RNA-seq reads at HERC2P3 location on chromosome 15. Data for all iPSC and NSC samples are group-autoscaled using Integrative Genome Viewer (IGV). **D.** mRNA levels of *ZNF560* in HC and SZ iPSC relative to passage numbers. *GAPDH* mRNA was used to normalize gene expression values and results were expressed relative to iPSC SZ1. **E.** Precision-Recall curve from RNA-seq data of ZNF560 from Figure 1J.

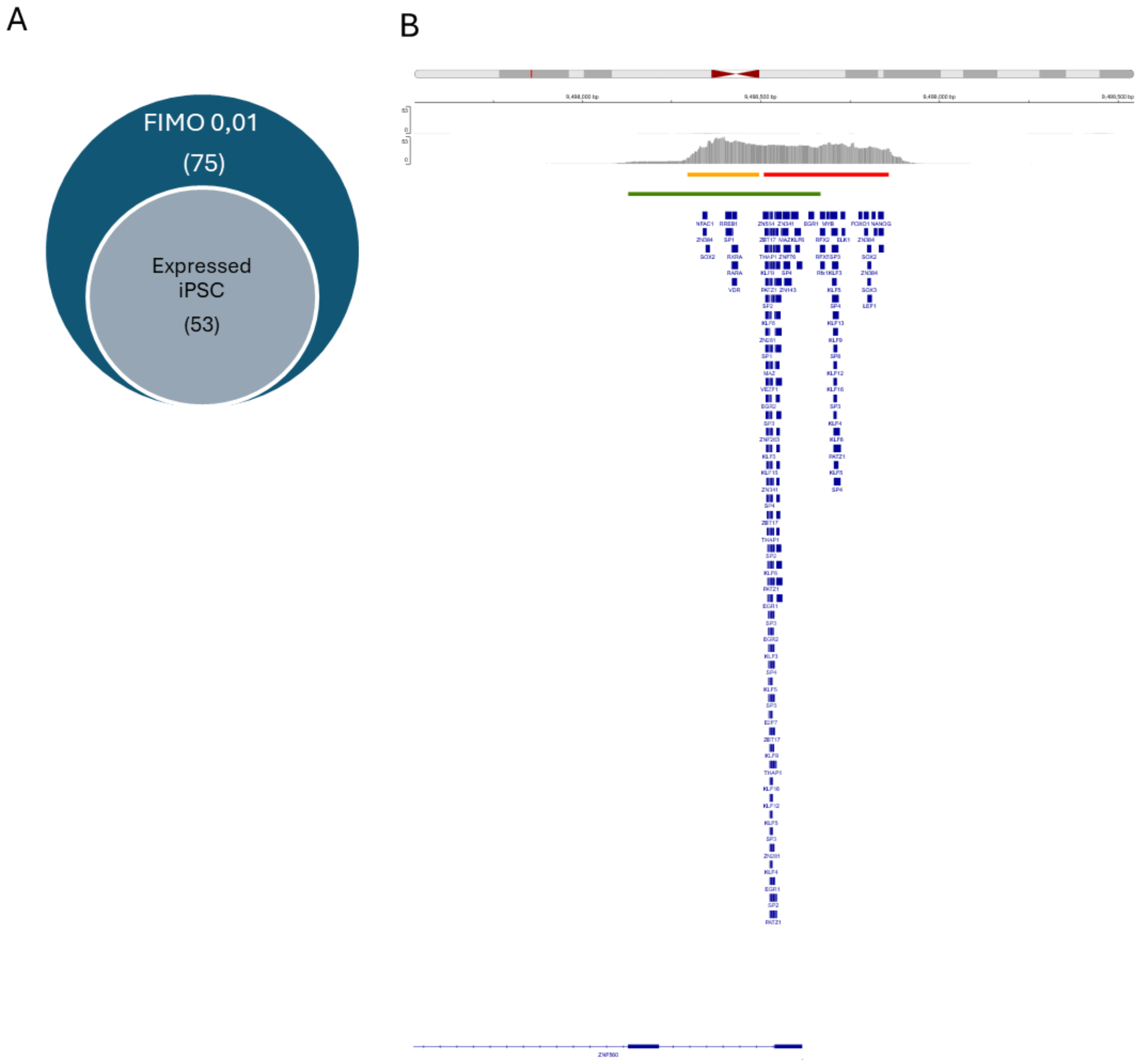


**Supplementary Figure 2. A.** Chart depicting the proportion of predicted ZNF560 promoter binding transcription factors that are expressed in iPSC relative to the total number of transcription factors with motifs identified by FIMO. **B.** Genomic visualization of the *ZNF560* gene promoter. Representative SZ-iPSC ATAC-seq signal track is shown in grey. Colored bars indicate presence of Cis-Regulatory Elements (CREs) obtained from the ENCODE database: red bar corresponds to promoter-like signatures, yellow bar to proximal enhancer-like signatures, and green bar to a CpG island. Blue bars indicate the location of predicted transcription factor motifs.

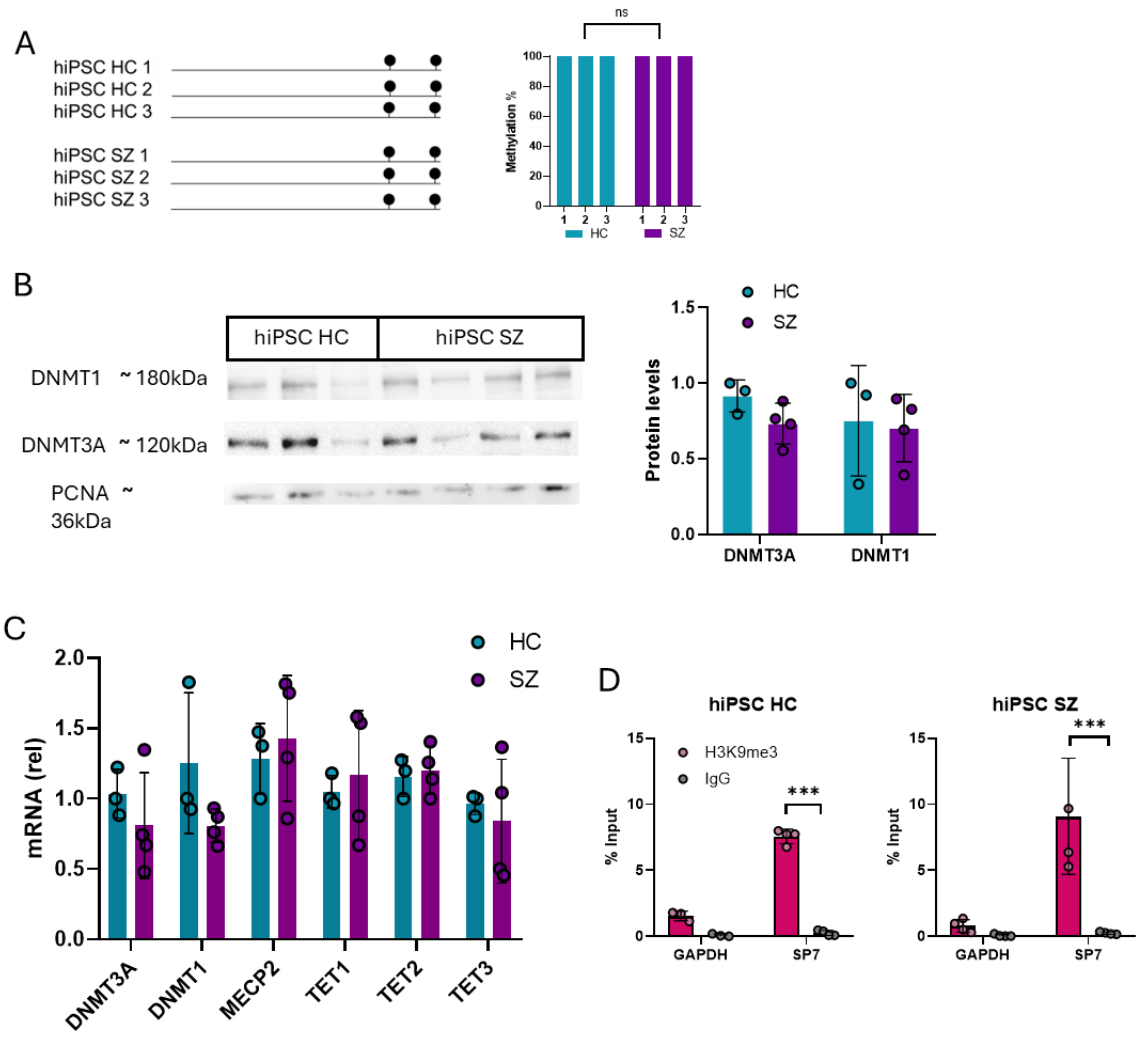


**Supplementary Figure 3**. **A.** Lollipop diagram of 5mC methylation at the *BGLAP* gene promoter. 10 clones were analyzed in each iPSC (3 HC and 4 SZ). **B**. Representative immunoblot analysis of DNMT1 (~180 kDa) and DNMT3A (~120 kDa) protein levels in 3 HC and 4 SZ iPSC. PCNA (~36 kDa) was used as a loading control. Quantification of DNMT3A and DNMT1 protein levels, normalized to PCNA, expressed relative to iPSC HC 1. Bars represent mean ± SD. Absence of significant differences was established by ANOVA. **C.** Relative mRNA expression levels of *DNMT3A*, *DNMT1*, *MECP2*, *TET1*, *TET2*, and *TET3* in iPSC (3 HC and 4 SZ) were determined by RT–qPCR. Expression was normalized to *GAPDH* mRNA levels and showed relative to iPSC HC1. No significant differences were established by ANOVA. **D.** ChIP–qPCR analysis showing enrichment (% input) of H3K9me3 at the promoter regions of *GAPDH* (active gene) and *SP7* (highly repressed gene) in iPSC (3 HC and 4 SZ). IgG was used as a non-specific precipitation control. Data is shown as mean ± SD, with ***p < 0.001, by 2-way ANOVA.

A

B

C

D

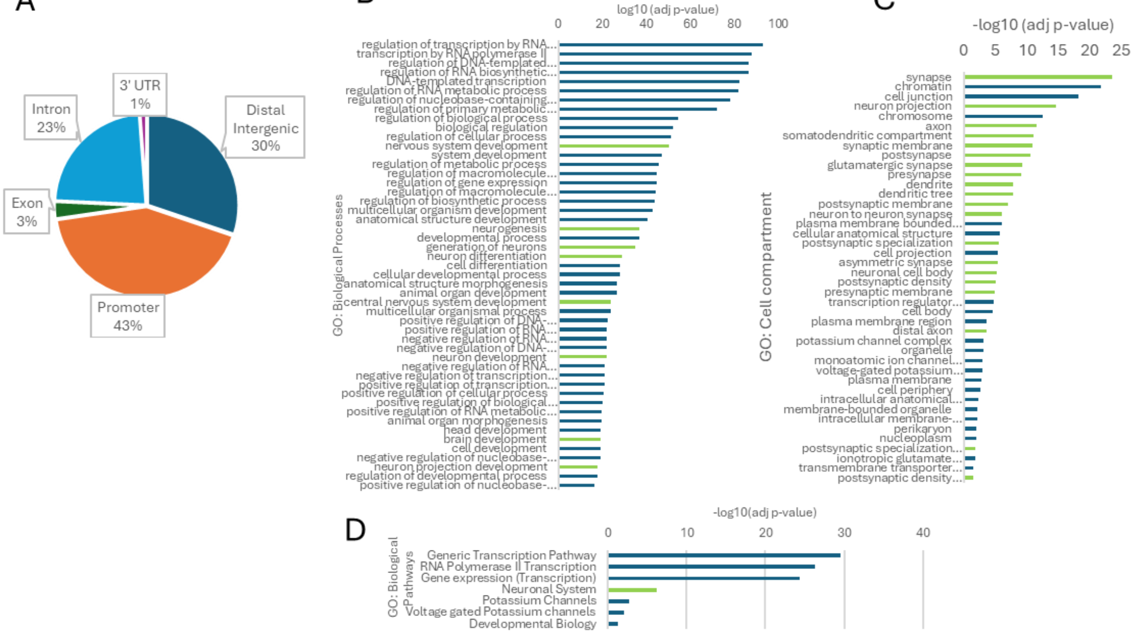


E

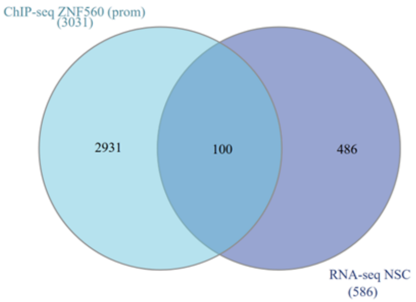


| source | term_name | term_id | -log10(adj. P-value) |
|---|---|---|---|
| GO:MF | sequence-specific double-stranded DNA binding | GO:1990837 | 95 |
| | DNA-binding transcription factor activity | GO:0003700 | 91 |
| | voltage-gated channel activity | GO:0022832 | 3 |
| | 3',5'-cyclic-GMP phosphodiesterase activity | GO:0047555 | 2 |
| | glutamate receptor activity | GO:0008066 | 1 |
| | phosphoric diester hydrolase activity | GO:0008081 | 1 |
| GO:BP | regulation of transcription by RNA polymerase II | GO:0006357 | 86 |
| | trans-synaptic signaling | GO:0099537 | 13 |
| | homophilic cell adhesion via plasma membrane adhesion molecules | GO:0007156 | 6 |
| | cellular response to endogenous stimulus | GO:0071495 | 6 |
| | cAMP-mediated signaling | GO:0019933 | 3 |
| | response to oxygen-containing compound | GO:1901700 | 2 |
| | cAMP catabolic process | GO:0006198 | 2 |
| | adenylate cyclase-modulating G protein-coupled receptor signaling pathway | GO:0007188 | 2 |
| | receptor localization to synapse | GO:0097120 | 2 |
| | vesicle-mediated transport in synapse | GO:0099003 | 1 |
| GO:CC | synapse | GO:0045202 | 24 |
| | chromatin | GO:0000785 | 19 |
| | transcription regulator complex | GO:0005667 | 5 |
| | monoatomic ion channel complex | GO:0034702 | 3 |
| | potassium channel complex | GO:0034705 | 3 |
| | synaptic cleft | GO:0043083 | 2 |

**Supplementary Figure 4. A.** The pie chart shows the genomic distribution of ZNF560 ChIP-seq peaks in the HEK293T cell line (GEO: GSM6047482). **B-D.** GO analysis of genes with ZNF560 binding at their promoters. Bar plots display the top-25 enriched GO terms associated with biological processes (B), cell compartments (C), and biological pathways (D). Terms related to neurodevelopment are highlighted in light green. **E.** Because ZNF560 has been proposed to function as a transcriptional repressor, we intersected the list of genes showing ZNF560 binding with the subset of genes downregulated in SZ-iPSC-derived SZ-NSC; the Venn diagram (left) reveals 100 shared genes. GO analysis of these shared genes (right) shows significant enrichment of neuronal-related terms.
